# Modulated TRPC1 expression predicts sensitivity of breast cancer to doxorubicin and magnetic field therapy: segue towards a precision medicine approach

**DOI:** 10.1101/2021.04.30.442085

**Authors:** Yee Kit Tai, Karen Ka Wing Chan, Charlene Hui Hua Fong, Sharanya Ramanan, Jasmine Lye Yee Yap, Jocelyn Naixin Yin, Yun Sheng Yip, Wei Ren Tan, Angele Pei Fern Koh, Nguan Soon Tan, Ching Wan Chan, Ruby Yun Ju Huang, Alfredo Franco-Obregón

## Abstract

**Background:** Chemotherapy is the mainstream treatment modality for invasive breast cancer. Nonetheless, chemotherapy-associated adverse events can result in a patient terminating treatment. We show that transient receptor potential channel 1 (TRPC1) expression level predicts breast cancer sensitivity to doxorubicin (DOX) and pulsed electromagnetic field (PEMF) therapies.

**Methods:** The effects of PEMFs were examined with respect to: **1**) the growth of MCF-7 cells *in vitro*; **2**) MCF-7 tumors implanted into a chicken chorioallantoic membrane (CAM) model and; **3**) patient-derived and MCF-7 breast cancer xenografts in mice.

Potential synergisms between DOX and PEMF therapies were examined in these model systems and under conditions of TRPC1 overexpression or silencing *in vitro*.

**Results:** PEMF exposure impaired the survival of MCF-7 cells, but not that of nonmalignant MCF10A breast cells. The effects of PEMF- and DOX-therapies synergized *in vitro* at compromising MCF-7 cell growth. Synergism could be corroborated *in vivo* with patient-derived xenograft mouse models, wherein PEMF exposure alone or in combination with DOX reduced tumor size. Stable overexpression of TRPC1 enhanced the vulnerability of MCF-7 cells to both DOX and PEMF exposure and promoted proliferation, whereas chronic DOX exposure reduced TRPC1 expression, induced chemoresistance, precluded response to PEMF exposure and mitigated proliferation. Markers of metastasis including *SLUG, SNAIL, VIMENTIN*, and *E-CADHERIN* as well as invasiveness were also positively correlated with TRPC1 channel expression.

**Conclusion:** The presented data supports a potential role of PEMF-therapy as an effective companion therapy to DOX-based chemotherapy for the treatment of breast cancers characterized by elevated TRPC1 expression levels.

## Background

Breast cancer is the leading cause of cancer-associated death for women worldwide (1). It is estimated that 1 in 8 women in the US will be diagnosed with invasive breast cancer within their lifetimes (2). And, although chemotherapy is the mainstream treatment modality for breast cancer, greater than 50% of women undertaking chemotherapy will experience at least one chemotherapy-related adverse event (3). An urgent need hence exists for companion therapies to improve chemotherapeutic outcome in hopes of mitigating associated adverse events and reducing treatment-related toxicities.

Doxorubicin (DOX) is the most widely used chemotherapeutic agent for breast and other cancers (3). The anticancer effects of DOX are attributed to its ability to inhibit DNA replication in actively-proliferating cancer cells (4) as well as to stimulate reactive oxygen species (ROS) production via a mechanism of redox cycling, causing oxidative damage to lipids, DNA, and proteins (3). The ensuing mitochondrial damage further accentuates DOX-dependent ROS production to exacerbate oxidative damage (4).

Brief exposure (10 min) to low amplitude (1 mT) pulsing magnetic fields (PEMFs) has been shown capable of stimulating mitochondrial respiration and ROS production (5), thereby promoting both *in vitro* (5) and *in vivo* (6) myogeneses via a process of magnetic mitohormesis. Obeying a mitohormetic mechanism of operation (7), brief and low amplitude PEMF exposure would produce low levels of ROS sufficient to instill mitochondrial survival adaptations, whereas exaggerated PEMF exposure might be expected to produce detrimental levels of oxidative stress that instead stymie cell survival. Importantly, the threshold for achieving an irreversibly damaging level of oxidative stress would depend on the basal metabolic rate and existing inflammatory status of the recipient cells. Accordingly, exposure to 3 mT PEMFs for one hour was previously shown to be cytotoxic to MCF-7 breast cancer cells, whereas the same exposure paradigm was tolerated by MCF10A nonmalignant breast cells (8).

Transient Receptor Potential Channel 1 (TRPC1) expression is necessary and sufficient to bestow PEMF-stimulated mitochondrial respiration and proliferation (9). Evidence of a TRPC1-mitochondrial axis exists with the findings that calcium entry modulates mitochondrial respiration (10), whereas mitochondrial ROS reciprocally modulates TRPC1 function (11). TRPC1-mediated calcium was hence proposed as an exploitable point of vulnerability to undermine cancer viability (12, 13) by commandeering the calcium/ROS-dependent cytotoxicity pathway (14, 15). TRPC1 and TRPM7 are the most abundantly expressed of all TRP channels (16), underscoring their well-documented physiological and clinical importance. Elevated expression levels of TRPC1, TRPC6, TRPM7, TRPM8, and TRPV6 are detected in human breast ductal adenocarcinoma (hBDA) cells (17), whereby the expressions of TRPC1, TRPM7, and TRPM8 were most closely correlated with proliferative deregulation and tumor growth, and TRPV6 was more strongly correlated in invasive breast cancers. On the other hand, in high histopathological grade breast cancers, TRPC1 expression was negatively correlated with invasiveness and chemoresistance (17). Conversely, DOX treatment has been shown to induce genotypic and phenotypic modifications that make cancer cells refractory to chemotherapy (18). While the chemotherapeutic agents, cisplatin and carboplatin, are capable of downregulating TRPC1 channel expression in ovarian cancer cell lines (19), the effect of DOX on TRPC1 channel expression in breast cancer is unexplored.

Given the reported capacity of PEMFs to target breast cancer cells (8), we hypothesized that DOX and PEMF treatments might synergize to undermine breast cancer growth. We provide relevant evidence *in vitro* and *in vivo* that combined DOX and PEMF treatments slow growth, augment apoptosis and enhance breast tumor resorption to a greater degree than either treatment alone. Evidence is also provided that TRPC1 expression level predicts breast cancer vulnerability to DOX and PEMF treatments. Overexpression of TRPC1 increased MCF-7 vulnerability to DOX and PEMF exposure, whereas silencing TRPC1 expression abrogated sensitivity to PEMFs and DOX and could be recapitulated with prolonged DOX-exposure. TRPC1 overexpression also increased MCF-7 cell proliferation and invasiveness, whereas TRPC1 downregulation suppressed proliferation, supporting a role for TRPC1 in tumorigenesis and providing a selective target for PEMF-intervention.

## Materials and methods

Full materials and methods are made available in Supplementary Information.

### Chick Chorioallantoic Membrane (CAM) Model

The chick chorioallantoic membrane (CAM) assay (20) was performed using fertilized Bovans Goldline Brown chicken eggs purchased from Chew’s Egg Farm Pte Ltd., Singapore. Briefly, eggs were placed horizontally in a 38.5°C humified chamber of 70% humidity for 3 days. On day 3, 3 to 4 ml of albumin was removed through a hole in the apex of the eggs using an 18G needle on a 5 ml syringe to lower the CAM. An oval 1 cm^2^ hole was then made on the center of the eggs and covered using a 1624W Tegaderm semi-permeable membrane. On day 7, the eggs were inoculated with 1.5 x 10^6^ MCF-7 cells resuspended in 50 ul of Matrigel (Sigma Aldrich) on the blood vessel of the CAM. Prior to the inoculation of the MCF-7 cells, the blood vessels closer to the CAM were gently perforated using a dry glass rod. The eggs were resealed using Tegaderm and left for another 2 days. The eggs were then exposed to PEMF stimulation on days 9, 10, and 11 for 1 h each day. Tumor weight was determined on Day 13.

### MCF-7 Breast Cancer Xenograft and Patient-Derived Xenograft (PDX) Model in NSG mice

NSG (NOD.Cg-Prkdc^scid^ Il2rg^tm1Wjl^/SzJ) mice, which lack human-specific cytokines and human leukocyte antigen (HLA) expression on stromal cells were used to host the breast cancer cell lines or patient-derived xenografts (PDX) (21). The NSG mice were purchased from Jackson’s laboratory and used at 8-10 weeks of age. Briefly, each female NSG mouse was implanted with a subcutaneous 60-day (0.36 mg) slow-release estradiol pellet (Innovative Research of America). Each patient tumor was equally divided into 5 chunks and implanted into the dorsal flank of 5 animals corresponding to the 5 different treatment groups. The tumors were allowed to grow for 3 weeks. For MCF-7, 1×10^6^ cells were counted and mixed in a 40-60% ratio with Matrigel Growth Factor (Bio Laboratories, Cat No. 354230). The cells are injected subcutaneously into the dorsal flank of the animals corresponding to the different treatment groups. The animals were given 20 mg/kg DOX intravenously and/or PEMFs stimulation for 1 h weekly for 5 weeks. At the end of the study, tumor volume was measured and isolated for apoptotic cell determination.

## Results

### PEMF exposure impairs breast cancer cell growth *in vitro* and *in vivo*

PEMF exposures at an amplitude of 3 mT administered for 3 consecutive days for 1 h per day were previously shown capable of impairing MCF-7 cell viability (Fig. 1A) (8). We extend these initial findings by showing that an analogous PEMF exposure paradigm reduced viable cell counts for both MCF-7 (∼34%; Fig. 1B) and MDA-MB-231 (∼19%; Fig. 1C) breast cancer cell lines, whereas MCF10A normal breast cell line subjected to the same PEMF paradigm did not exhibit a change in cell number (Fig. 1D) relative to unexposed (0 mT) cells. Moreover, stronger PEMF exposures (5 mT) were ineffective at killing MCF-7 and MDA-MB-231 breast cancer cells (Fig. 1B, 1C & 1D). The long-term effects of magnetic field exposure were examined in the context of colony formation (22), whereby MCF-7 cells were plated at clonal density and exposed to 3 mT PEMFs for 10 successive days (Fig. 1E). Colony number (Fig. 1F) and size (Fig. 1G) were reduced relative to unexposed (0 mT) cultures, consistent with the ability of PEMFs to reduce MCF-7 cell number (Fig. 1B).

**Figure 1.**
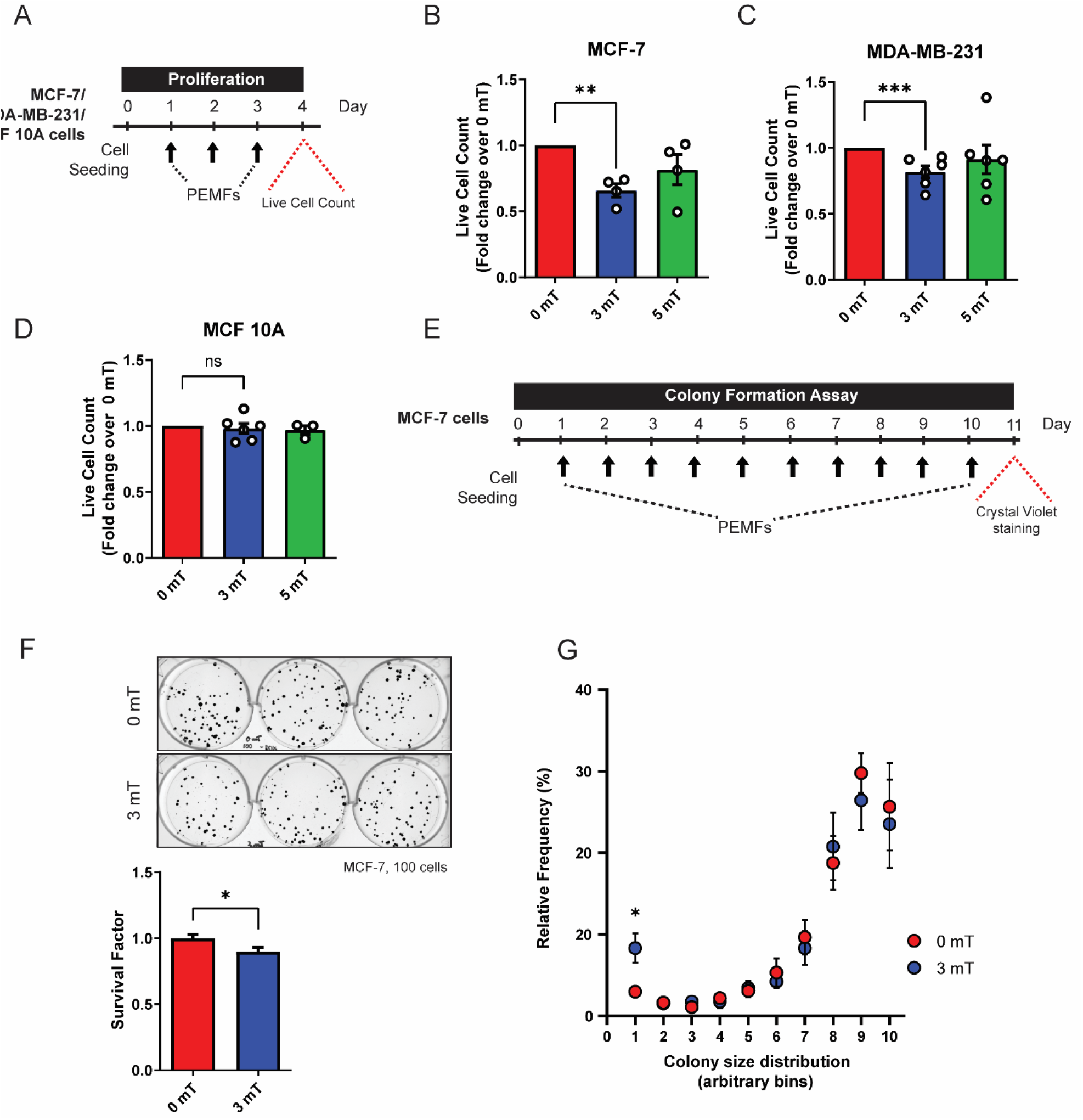
PEMF exposure inhibits cancer cell growth *in vitro*. **A)** Schematic of PEMF exposure schedule for live cell quantification. **B, C, and D)** Live cell counts using Trypan Blue exclusion assay for MCF-7, MDA-MB-231 and MCF10A cells. Cells were exposed to 0 mT, 3 mT, or 5 mT PEMF for 1 h each day for 3 consecutive days before cell count was performed. **E)** Schematic of colony formation assay schedule for MCF-7 cells analyzed after 10 daily PEMFs exposure at 3 mT for 1 h. **F)** Representative images of MCF-7 cell colony formation assay. Cells were seeded at 100 cells per well. Survival factor represents the number of surviving colonies per 100 cells and presented as fold change over 0 mT. **G)** Relative frequency of colonies (in Figure F) according to their arbitrary sizes binned into 10 bins per total number of cells. All experiments were of at least 3 independent replicates performed with ^*\**^*p* < 0.05, ^*\*\**^*p* < 0.01, ^*\*\*\**^*p* <0.001. The error bars represent the standard error of the means.

To more closely approximate the *in vivo* scenario, the chicken chorioallantoic membrane (CAM) model was employed to explore the capability of PEMF exposure to modulate tumor growth (20). In this animal model, the absence of an immune system during early chick development allows for the stable growth of breast cancer tumors. MCF-7-derived tumors were implanted into 7-day old eggs and commencing the following day exposed to 3 mT for 1 h per day for 3 consecutive days (Fig. 2A). Tumor xenografts exposed to 3 mT showed a substantial loss in tumor weight of ∼50% compared to unexposed (0 mT) tumors (Fig. 2B).

**Figure 2.**
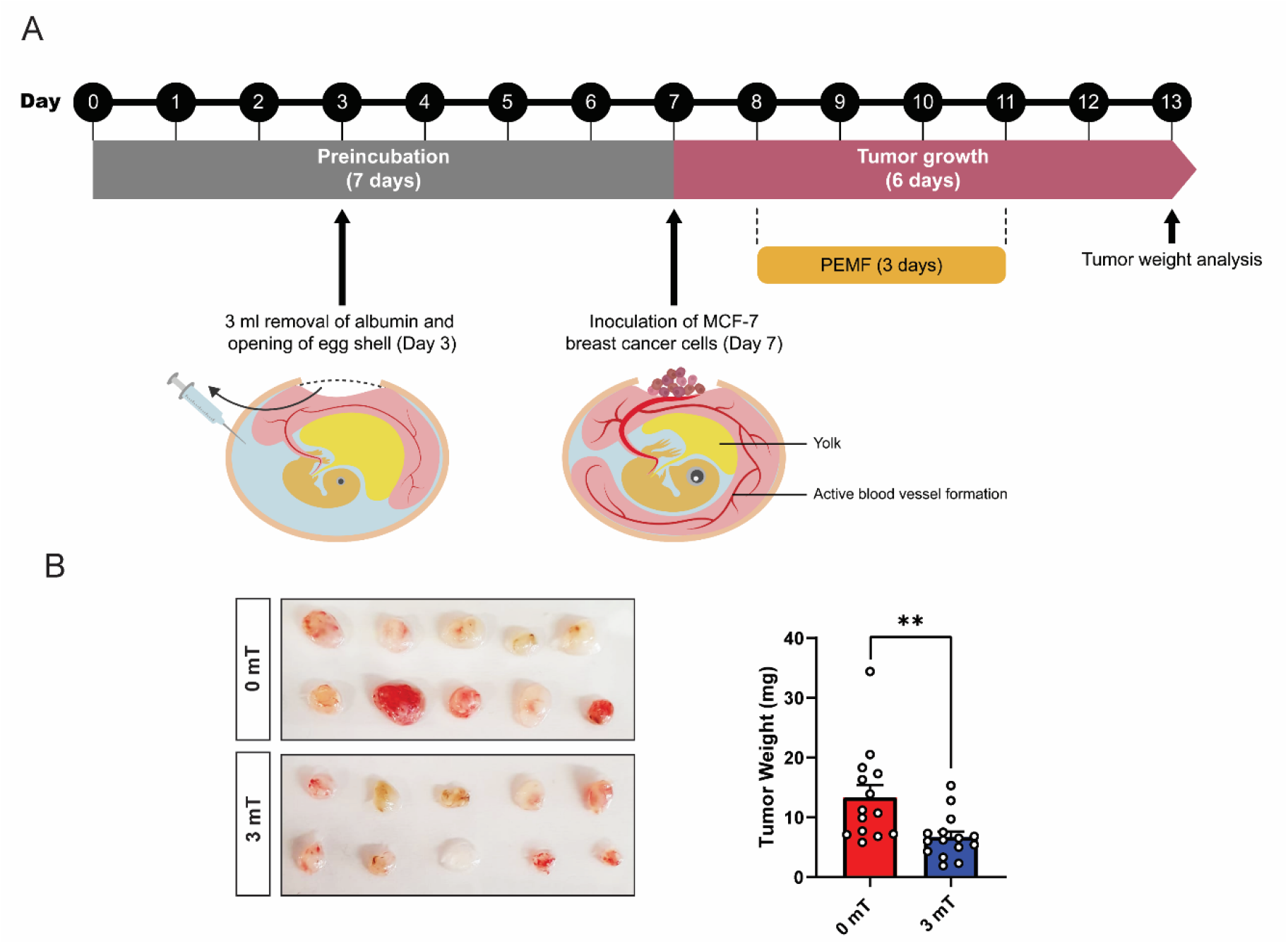
PEMF inhibits breast tumor growth *in vivo*. **A)** Schematic of the PEMF exposure paradigm used on the CAM model for MCF-7 breast tumor xenografts. MCF-7 tumors were inoculated onto CAM on day 7. The eggs were exposed to 3 mT for 1 h for 3 successive days and left to grow for another 3 days before weight analysis. **B)** Images showing the size of MCF-7 tumors and the corresponding bar chart represents the pooled tumor weight (mg). Experiments were repeated twice with ^*\*\**^*p* < 0.01 of at least 14 independent eggs. The error bars represent the standard error of the mean.

### PEMF exposure increases the vulnerability of cancer cells to doxorubicin

We examined whether PEMF exposure modulates chemotherapeutic efficacy in breast cancer cells. Proliferative and colony-forming capacities were ascertained in response to 3 consecutive days of PEMF exposure in combination with 100 nM DOX, the reported DOX IC_50_ in MCF-7 cells (23), administered on the third day (Fig. 3A). Exposure of MCF-7 cell cultures to PEMFs alone for 1 h per day for 3 consecutive days reduced the cellular DNA content by ∼20% relative to unexposed control cultures (Fig. 3B; solid red, 0 mT; solid blue, 3 mT). DOX treatment alone reduced DNA content by ∼35% (hatched red). The preconditioning of MCF-7 cells with two days of PEMF exposure (1 h/day) accentuated DOX-cytotoxicity by an additional ∼18% (Fig. 3B; hatched blue). The effects of PEMF exposure during prolonged DOX treatment were ascertained by reducing the concentrations of DOX to 10 nM (5-fold) or 20 nM (10-fold), followed by colony analysis. MCF-7 cultures were administered DOX on days 1, 4, and 7 in conjunction with daily PEMF exposure for 10 days (Fig. 3C). PEMF exposure in combination with chronic DOX administration reduced MCF-7 colony formation (Fig. 3D; bottom) more than DOX treatment alone (Fig. 3D; top). For low-density cultures, colony numbers (survival factor) were reduced by 10% and 27% in response to PEMF (Fig. 3E; solid blue) or DOX (Fig. 3E; hatched red) treatments alone, respectively, whereas combining PEMF and DOX treatments reduced colony number by 40% (Fig. 3E; hatched blue), relative to unexposed 0 mT cultures (Fig. 3E; solid red). The combination of treatments also increased and decreased the size of the remaining smaller and larger colonies, respectively (Fig. 3F; bottom), consistent with a slowing of cancer growth rate at the higher DOX concentration.

**Figure 3.**
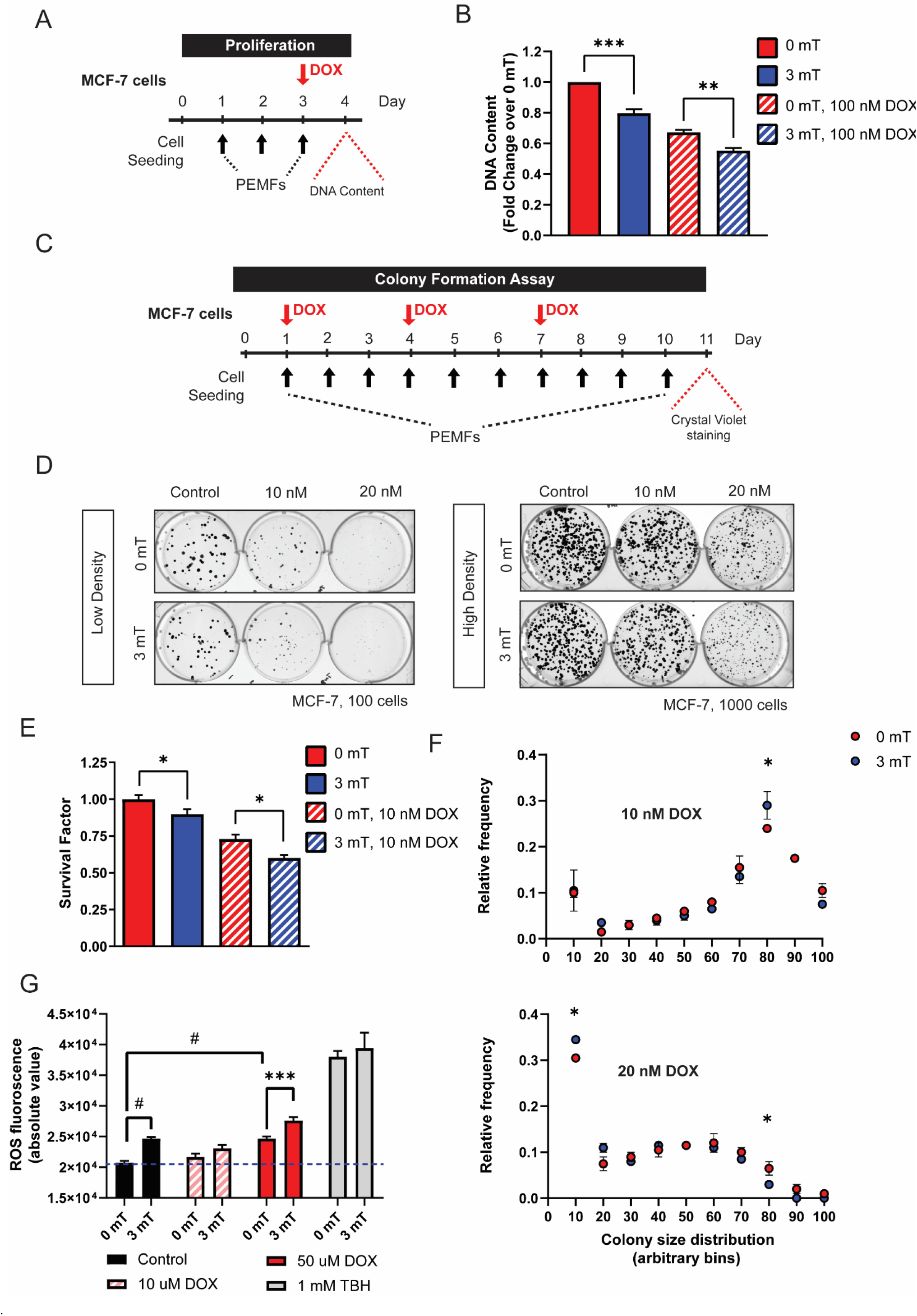
PEMF enhances the vulnerability of cancer cells to doxorubicin *in vitro*. **A)** Schematic of 3 mT PEMF and DOX treatment on MCF-7 cells for Cyquant DNA quantification in 96-well plate format. Cells were exposed to 3 mT for 1 h daily for 3 successive days. DOX (100 nM) was treated on the final day 1 h before the last PEMF exposure. Cellular DNA content was measured 24 h after the last PEMF exposure. **B)** Bar chart shows the pooled data for Cyquant DNA content in fold change 24 h post-DOX and PEMF treatments. **C)** Schematic of colony formation assay for MCF-7 cells treated with DOX and 3 mT exposure for 10 days. **D)** Representative images of colony formation of MCF-7 cells showing dose-response with 10 and 20 nM DOX, with 0 or 3 mT PEMFs. Cells were seeded at 100 or 1000 cells per well to show the combined effect of DOX and PEMFs. **E)** The corresponding bar chart shows the colony survival factor in fold-change (over 0 mT, solid red) for 10 nM DOX under low-density condition. **F)** Relative frequency of colonies per total number of cells binned according to colony size for treatment of 10 nM (top) or 20 nM (bottom) DOX under high-cell density condition, with 0 mT and 3 mT marked by red and blue dots, respectively. **G)** Bar charts show the absolute ROS fluorescence of DCH_2_FDA on MCF-7 cells. Cells were incubated in DCH_2_FDA for 30 min before washing and replacement with media containing DOX (hatched pink and red) or TBH (grey). Cells were exposed to 3 mT for 10 min and immediately thereafter ROS fluorescence measurement after 20 min. The mean ROS fluorescence presented is an average of 8 technical replicates. All data presented were performed with at least 3 independent experiments with ^*\**^*p* < 0.05, ^*\*\**^*p* < 0.01, ^*\*\*\**^*p* < 0.001 ^*#*^*p* < 0.0001. The error bars represent the standard error of the mean.

PEMF exposure stimulates ROS production in cancer (24, 25) and non-cancer (5, 26) cells. Underlying this response is a magnetically-sensitive, TRPC1-mediated calcium entry pathway modulating mitochondrial respiration (5, 27). On the other hand, DOX increases cytoplasmic and mitochondrial ROS by disrupting mitochondrial redox cycling and function (28). Lone PEMF exposure of MCF-7 cells (Fig. 3G; solid black) increased ROS levels by ∼19% over baseline (0 mT). By comparison, tert-Butyl hydroperoxide (TBH; 1 mM), a cytoplasmic pro-oxidant, increased ROS levels by ∼83% that furthermore, quenched a subsequent response to PEMFs (Fig. 3G; grey). The acute administration of 10 or 50 uM DOX increased ROS levels by ∼7% and ∼12%, respectively, that was further augmented by PEMF exposure by ∼4% (Fig. 3G; hatched pink) and ∼19% (Fig. 3G; red), respectively. PEMF and DOX (50 uM) treatments hence synergistically act to raise ROS levels in naïve MCF-7 cells.

### PEMF exposure enhances the vulnerability of breast tumors to DOX *in vivo*

Patient-derived breast tumor xenografts (PDX) were engrafted into immunocompromised NSG mice and allowed to grow for 3 weeks before once-weekly administration of 20 mg/kg DOX and/or exposure to PEMFs (3 mT) for 1 h per week. After 5 weeks of the indicated treatment, tumors were measured and analyzed by flow cytometry (Fig. 4A). Whereas untreated (control) tumors showed a progressive increase in volume of 87% from their initial values, the PEMF and DOX interventions instead reduced tumor volume by −55.4% and −69.8%, respectively (Fig. 4B). By contrast, the incidence of apoptotic cells increased by +1.61%, +8.8%, and +17.9% in tumors isolated from control, PEMF- and DOX-treated mice, respectively (Fig. 4C and 4D). Potential synergisms between DOX and PEMF interventions were ascertained with two paradigms: (1) once weekly PEMF treatment for 2 weeks followed by 3 weeks of DOX treatment alone and; 2) simultaneous weekly PEMF and DOX treatments for 5 weeks. Amongst all the test conditions, paradigm 1 (Fig. 4B, green) produced the greatest reductions in tumor volume (−80.9%) and increases in apoptotic cells (+44.6%) (Fig. 4D, green), wherein tumor resorption (Fig. 4B, green) was statistically different from DOX treatment alone (brown), but not from paradigm 2 (yellow). The livers from the PEMF- and DOX-treated mice showed little signs of collateral apoptosis (Fig. 4D, black), demonstrating cytotoxic specificity for malignant tissues by the employed PEMF paradigm.

**Figure 4.**
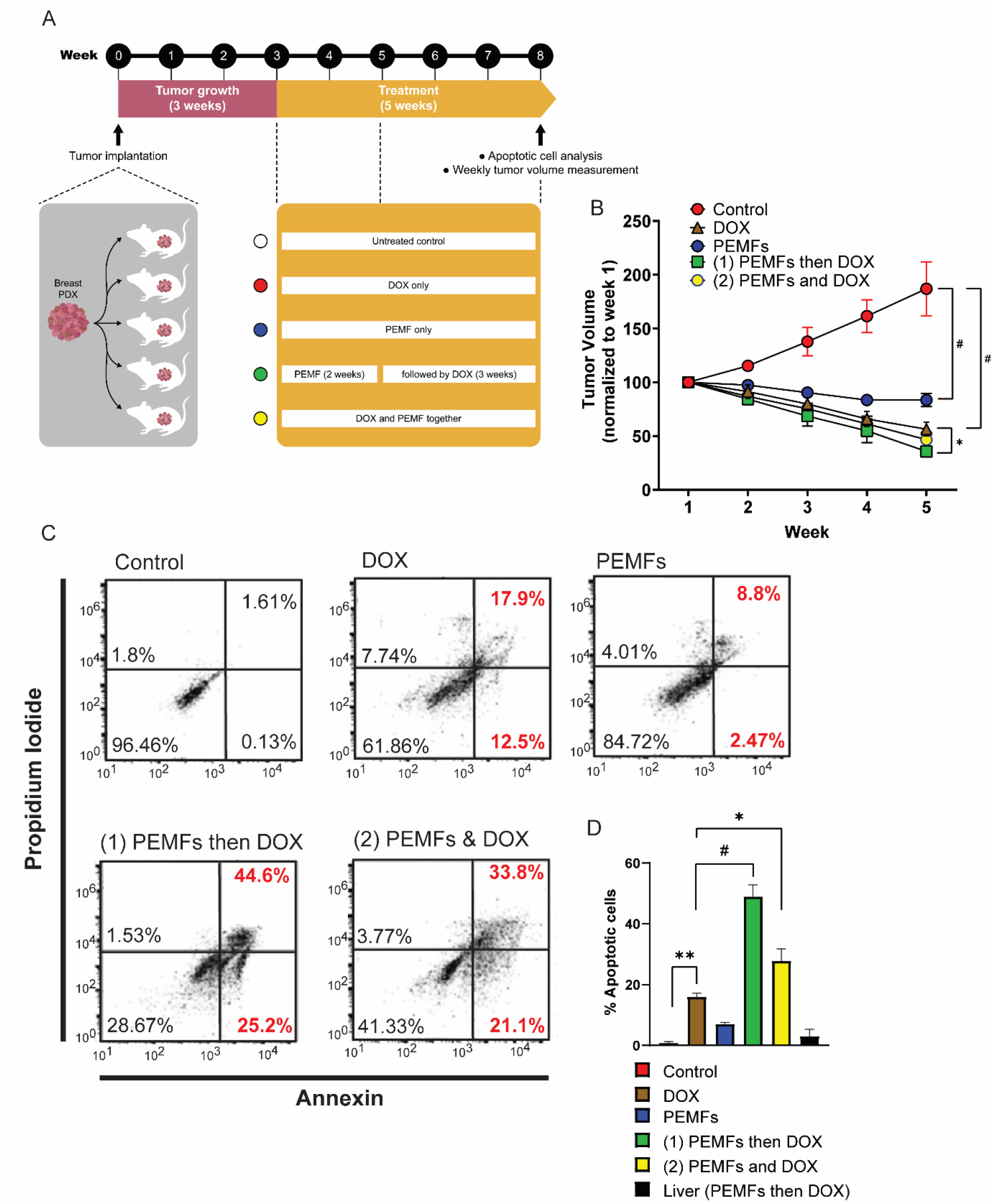
PEMFs synergize with DOX to inhibit tumor growth *in vivo*. **A)** Schematic of PEMF and DOX exposures weekly on patient-derived tumor xenograft in mice. Implanted tumors were allowed to grow for 3 weeks before the initiation of DOX and/or PEMF treatment. Tumor volume was measured every week while apoptotic cell determination was performed at the end of the study. Each data point represents the mean from 5 experimental runs derived from the tumors obtained from 5 patients, each of which was equally divided amongst the 5 treatment groups. **B)** Point graph showing the pooled data of tumor volume (mm^3^), measured for 5 consecutive weeks. **C)** Representative scatter dot-plots showing cell population of dissociated tumors sorted based on annexin and propidium iodide staining. **D)** Bar chart represents pooled data of apoptotic cell percentages analyzed using flow cytometry. N = 5 mice, with ^*\**^*p* < 0.05, ^*\*\**^*p* < 0.01, and ^*#*^*p* < 0.0001. The error bars are expressed as the standard error of the mean.

We also examined the effects of PEMF exposure on MDA-MB-231 and MCF-7 breast cancer cells engrafted into NSG mice. MDA-MB-231 tumors from NSG mice exposed once (3 mT x 1) or twice (3 mT x 2) to PEMFs exhibited increases in apoptosis of +11% and +34% respectively, over baseline (0 mT) (Supplementary Fig. 1A and 1B). Livers harvested from these mice similarly did not show any significant increase in apoptosis (Supplementary Fig. 1C). MCF-7 xenografts were also subjected to the same PEMF/DOX paradigm previously employed in Figure 4 (Supplementary Fig. 2A). Again, PEMF and DOX treatments synergized to promote cancer cytotoxicity, achieving +24% and +33% apoptosis (early plus late) for tumors subjected to paradigms 1 (PEMF then DOX) and 2 (PEMF and DOX), respectively (Supplementary Fig. 2B). The percentages of apoptosis obtained from paradigms 1 and 2 was greater than those achieved with lone DOX (+14%), PEMF (+8%), or baseline (0 mT) (+0.3%) treatments. Although these responses were more modest than previously obtained in the PDX mouse trial (Fig. 4), synergism between DOX and PEMF treatments in undermining *in vivo* cancer growth remained evident.

### Chronic DOX exposure reduces TRPC1 expression resulting in DOX-chemoresistance

PEMF exposure enhances TRPC1-mediated calcium entry and consequent engagement of the calcineurin-NFAT signaling axis involved in cellular homeostasis (5). In certain cancers, however, TRP hyperactivity may overwhelm NFAT-mediated (29) calcium and mitochondrial cellular homeostatic mechanisms (30), negatively selecting against cancer cells with inherently high TRP channel expression. To elucidate potential commonalities in mechanisms of action, we investigated TRPC1 channel expression levels in MCF-7 cells surviving either chronic PEMF and/or DOX exposures. MCF-7 cells were exposed to PEMFs for 10 consecutive days with the renewal of a sub-IC_50_ dose of DOX (20 nM) on days 1, 4, and 7 (Fig. 5A). Chronic DOX treatment alone (Fig. 5B; hatched red) reduced TRPC1 protein expression to 53% of control levels (solid red) and could be further reduced to 37% (hatched blue) of control levels with combined DOX and PEMF chronic treatment. By contrast, daily PEMF exposure alone did not reduce TRPC1 expression (solid blue). Furthermore, growth under chronic and progressive DOX treatment (≤ 96 nM) produced a stable DOX-resistant MCF-7 cell line (MCF-7/ADR) exhibiting attenuated TRPC1 expression (−36%) (Fig. 5C; hatched black) and proliferation (−90%) (Fig. 5D; hatched black). When serially passaged (>5 times) in the absence of DOX, however, MCF-7/ADR cells partially regained proliferative capacity and *TRPC1* expression (Fig. 5D, yellow). Therefore, chronic DOX exposure at the predetermined cytotoxic dose (23) was shown capable of negatively selecting against breast cancer cells with innately elevated TRPC1 expression to produce cellular progeny elaborating depressed TRPC1 expression, proliferative capacity and chemosensitivity. These findings reveal a relationship between DOX sensitivity and TRPC1 expression levels, aligning with previous findings that DOX targets proliferating cells (31) and that TRPC1 promotes proliferation (5).

**Figure 5.**
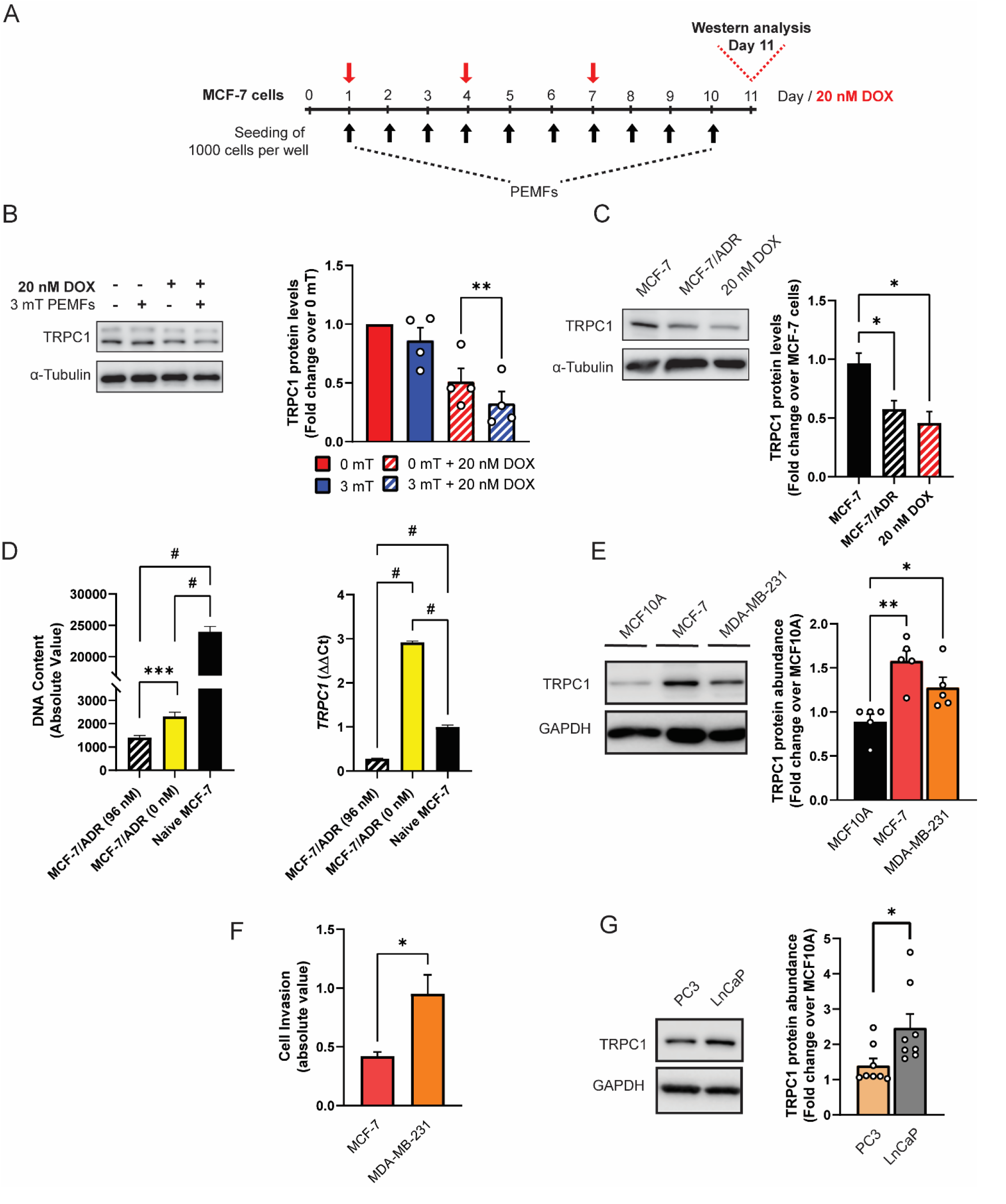
DOX-chemoresistance and metastatic status are associated with TRPC1 channel downregulation. **A)** PEMF and DOX (red arrow) treatment paradigm used for TRPC1 protein analysis. **B)** Representative western blot showing the changes in TRPC1 levels after PEMF and DOX treatment on day 11. The corresponding bar chart represents pooled data of TRPC1 protein levels normalized to unexposed 0 mT. Cells treated with DOX are represented by the hatched bars, with either 0 or 3 mT PEMFs. **C)** Representative western blot showing the relative fold change of TRPC1 protein in MCF-7/ADR (96 nM DOX; hatched black) and 11-day 20 nM DOX-treated MCF-7 cells (red) relative to naïve MCF-7 cells (solid black). Naïve MCF-7 and MCF-7/ADR cells were grown in culture for 3 days before western analysis. **D)** Cell growth comparison (72 h post-seeding) and *TRPC1* transcript levels between MCF-7/ADR (96 nM), MCF-7/ADR (0 nM) and naïve MCF-7 cells. MCF-7/ADR (96 nM) cells were generated using progressive DOX treatment up to 96 nM (hatched black). MCF-7/ADR (0 nM) corresponds to the cell whereby MCF7/ADR (96 nM) cells were serially passaged in the absence of DOX to give rise to MCF-7/ADR (0 nM) (yellow). **E)** Western analysis showing the relative expression of TRPC1 channel protein in non-malignant (MCF10A) and malignant breast cancer cells (MCF-7 and MDA-MB-231) after 48 h of growth under standard conditions. The corresponding bar chart shows the pooled data in fold change of TRPC1 expression normalized to MCF10A. **F)** Cell invasion comparison between breast cancer malignant cell lines, MCF-7 and MDA-MB-231 cells. Cells were seeded at high density and analyzed 48 h post-seeding. **G)** TRPC1 protein expression in metastatic prostate cancer cell lines PC3 and LnCaP (invasive status: PC3 > LnCaP). All results presented are of at least 3 independent experiments with ^*^*p* < 0.05, ^**^*p* < 0.01^***^, *p* < 0.001, and ^*#*^*p* < 0.0001. The error bars represent the standard error of the mean.

Basal TRPC1 channel expression was next correlated to malignancy status. MCF-7 and MDA-MB-231 breast cancer cells exhibited higher relative abundances of TRPC1 protein than the non-malignant MCF10A breast cells. Specifically, we found MCF-7 cells showed greater TRPC1 expression than the more malignant MDA-MB-231 cells (32) (Fig. 5E). Notably, MDA-MB-231 cells were also more invasive than MCF-7 cells as determined by their ability to infiltrate through a basement membrane (Fig. 5F), in agreement with published studies (32, 33). Higher TRPC1 channel expression was also observed in the less invasive LNCaP prostate cancer cell line relative to the highly metastatic PC3 prostate cancer cell line (34) (Fig. 5G), recapitulating the correlation between TRPC1 channel expression and metastatic status.

### TRPC1 overexpression enhances MCF-7 proliferation and EMT, but attenuates migratory capacity

A potential interplay between TRPC1 channel expression and proliferative, migratory capacities and epithelial-mesenchymal transition (EMT) indices was investigated. A TRPC1-GFP fusion protein overexpressing MCF-7 cell line (MCF-7/TRPC1) was generated and validated by western (Fig. 6A), qPCR (Fig. 6B), and immunofluorescence (Fig. 6C) analyses. As a relevant negative control, a stable cell line expressing the GFP vector was also generated. The GFP-TRPC1 fusion protein was highly expressed in the MCF-7/TRPC1 cells (Fig. 6A), exhibiting a 12-fold increase in *TRPC1* transcript levels compared to vector cells (Fig. 6B). Enhanced fusion protein expression was also verified using fluorescence imaging (Fig. 6C). MCF-7/TRPC1 (green) exhibited enhanced proliferation compared to vector-transfected cells (black) (Fig. 6D) and was corroborated by the elevated protein expression of Cyclin D1 (Fig. 6E). On the other hand, MCF-7/TRPC1 cells migrated more slowly (Fig. 6F; bottom) than vector control cells (Fig. 6F; top), manifested as a delayed closure of an introduced gap (Fig. 6G). TRPC1 overexpression also increased the gene expression of the EMT transcriptional activators, *SLUG*, and *SNAIL* (Fig. 6H), responsible for the metastatic reprogramming (35). Consistent with published findings, *SLUG* activation increased *VIMENTIN* transcript (Fig. 6H) and protein (Fig. 6I) levels (35), concomitant with decreases in E-cadherin transcript (Fig. 6H) and protein (Fig. 6J) levels, in accordance with transcriptional inhibition of E-Cadherin by Slug and Snail (36) and reported E-cadherin modulation in breast cancer tumors (37) and cells (38). Conversely, *TRPC1* silencing by dsiRNA transfection resulted in the downregulation of *SLUG* and *VIMENTIN* transcripts, with corresponding increases in *E-CADHERIN* transcripts, while *SNAIL* levels remained unchanged (Fig. 6K). The dsiRNA silencing of TRPC1 in naïve MCF-7 cells also reduced basal proliferation relative to scrambled RNA-transfected cells (Fig. 6L), corroborating the role of TRPC1 as a proliferation modulator and providing further evidence for TRPC1 involvement in breast cancer metastatic reprogramming.

**Figure 6.**
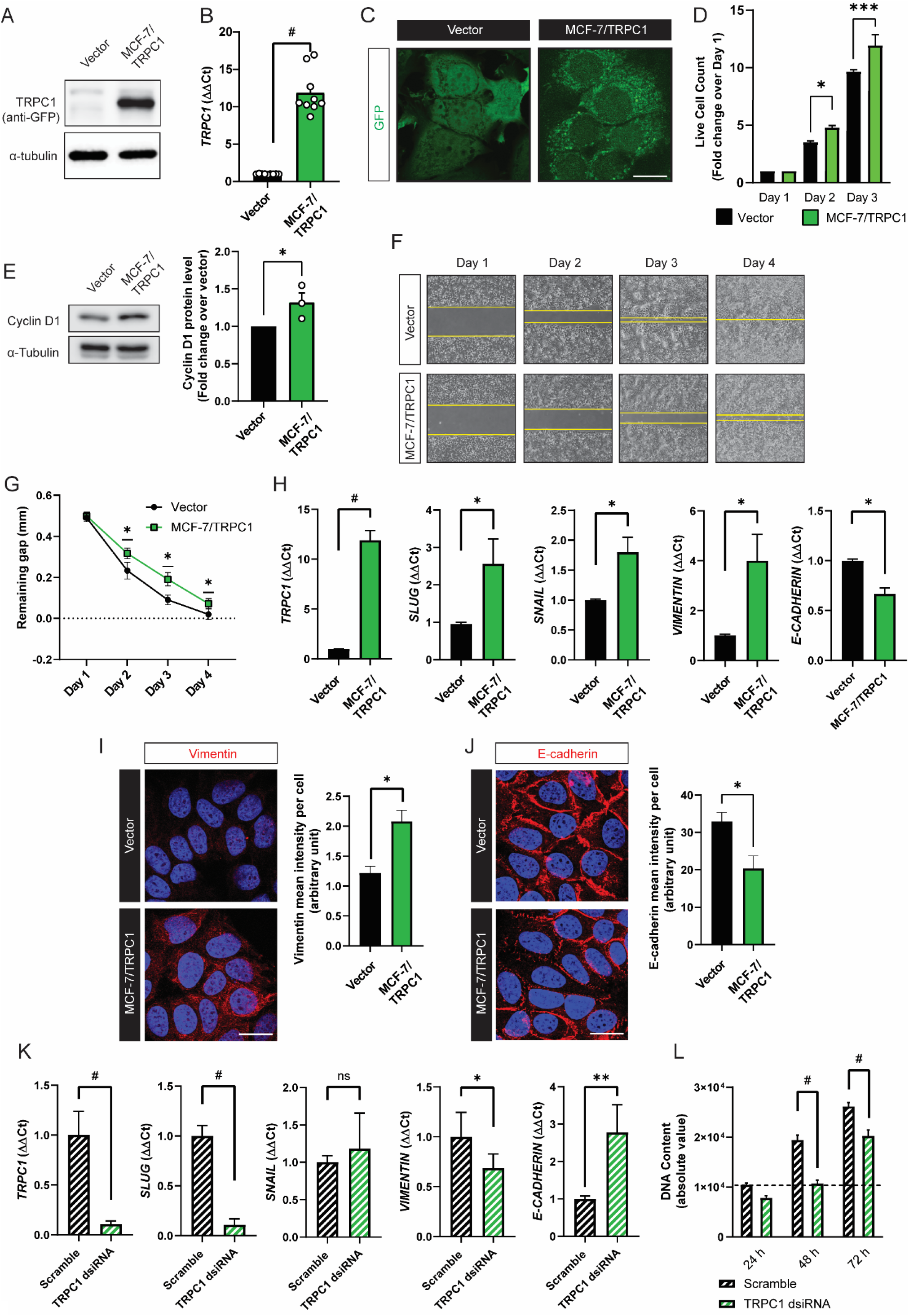
Characterization of TRPC1 overexpressing MCF-7 cell line. **A)** Western analysis showing the overexpression of GFP-TRPC1 in TRPC1 cells, stained using anti-GFP antibody. **B)** Bar chart shows ΔΔCt fold change of *TRPC1* transcript in MCF-7/TRPC1 cells (green) and vector-transfected cells (black). **C)** Fluorescence images showing GFP and GFP-TRPC1 in vector and MCF-7/TRPC1 cells, respectively. Scale bar = 10 µm. **D)** Bar chart showing live cell counts of stable cells over 3 days. **E)** Western analysis showing cyclin D1 protein levels 24 h post-seeding. **F)** Representative images of the migration assay over 4 days. Stable cells were seeded at high density one day before the removal of the insert to create a 0.5 mm gap. **G)** The corresponding line chart shows the remaining gap over 4 days for the migration assay. **H)** Transcript levels of *TRPC1, SLUG, SNAIL, VIMENTIN*, and *E-CADHERIN* in vector and MCF-7/TRPC1 cells. **I) and J)** Representative confocal images of vector and MCF-7/TRPC1 cells stained for Vimentin and E-Cadherin, with the corresponding bar charts showing mean intensity per cell, measured using absolute fluorescence intensity normalized to the total number of nuclei per view. **K)** Bar charts showing the transcript expression of *TRPC1, SLUG, SNAIL, VIMENTIN*, and *E-CADHERIN* in scrambled- and *TRPC1*-silenced cells. **L)** Examination of cell proliferation over 3 days using Cyquant DNA content analysis on *TRPC1*-silenced cells in relative to scramble RNA-transfected cells. *TRPC1* silencing was achieved using 2 independent dsiRNAs and the bar charts show the pooled data from the respective experiments. All results were of at least 3 independent experiments with ^*^ *p* < 0.05, ^**^ *p* < 0.01, ^***^ *p* < 0.001, ^#^ *p* < 0.0001. The error bars represent the standard error of the mean.

### PEMF exposure slows migration and increases invasiveness in TRPC1-overexpressing breast cancer cells

PEMF exposure (3 mT) further decelerated the migration of MCF-7/TRPC1 cells compared to unexposed (0 mT) MCF-7/TRPC1 cells (Fig. 7A, B). By contrast, the vector cells were insensitive to PEMF exposure (Fig. 7A, C). Invasiveness was ascertained by examining the ability of cells to break down, penetrate, and transverse a basement membrane-coated insert (Fig. 7D). According to this criteria, MCF-7/TRPC1 cells exhibited an invasive phenotype (Fig. 7D; green), comparable in magnitude to TGFβ-stimulated control cells (grey) and exceeding that of unstimulated control cells (black). PEMF exposure, attenuated the invasive capacity of MCF-7/TRPC1 cells (hatched green), but not of TGFβ-stimulated cells (hatched grey). As the invasiveness of MCF-7/TRPC1 was accompanied by an increase in the number of non-invading cells on the upper side of the culture insert (Fig. 7E), a causal relationship may exist between proliferation rate and invasive capacity as ascertained by this assay that initially may seem paradoxical given that TRPC1 overexpression (shown to augment proliferation; Fig. 6D and 6E) promotes invasiveness while slowing migration. On the other hand, the finding that PEMF exposure reduces invasiveness by attenuating both cell proliferation and migratory capacity is internally consistent and clinically exploitable. Agreeing with demonstrated transcriptional inhibition of E-cadherin in response to elevations in *SNAIL* and *SLUG* (Fig. 6H, J), E-cadherin protein levels were found to be reduced in MCF-7/TRPC1 cells (Fig. 7F). However, E-cadherin levels were unchanged by PEMF exposure (Fig. 7F), possibly reflecting an offsetting combination of PEMF-mediated TRPC1 downregulation (Fig. 5B), augmenting E-cadherin levels (Fig. 6K), and a PEMF-induced slowing of cell migration (Fig. 7A, B), reinstating E-cadherin levels (39).

**Figure 7.**
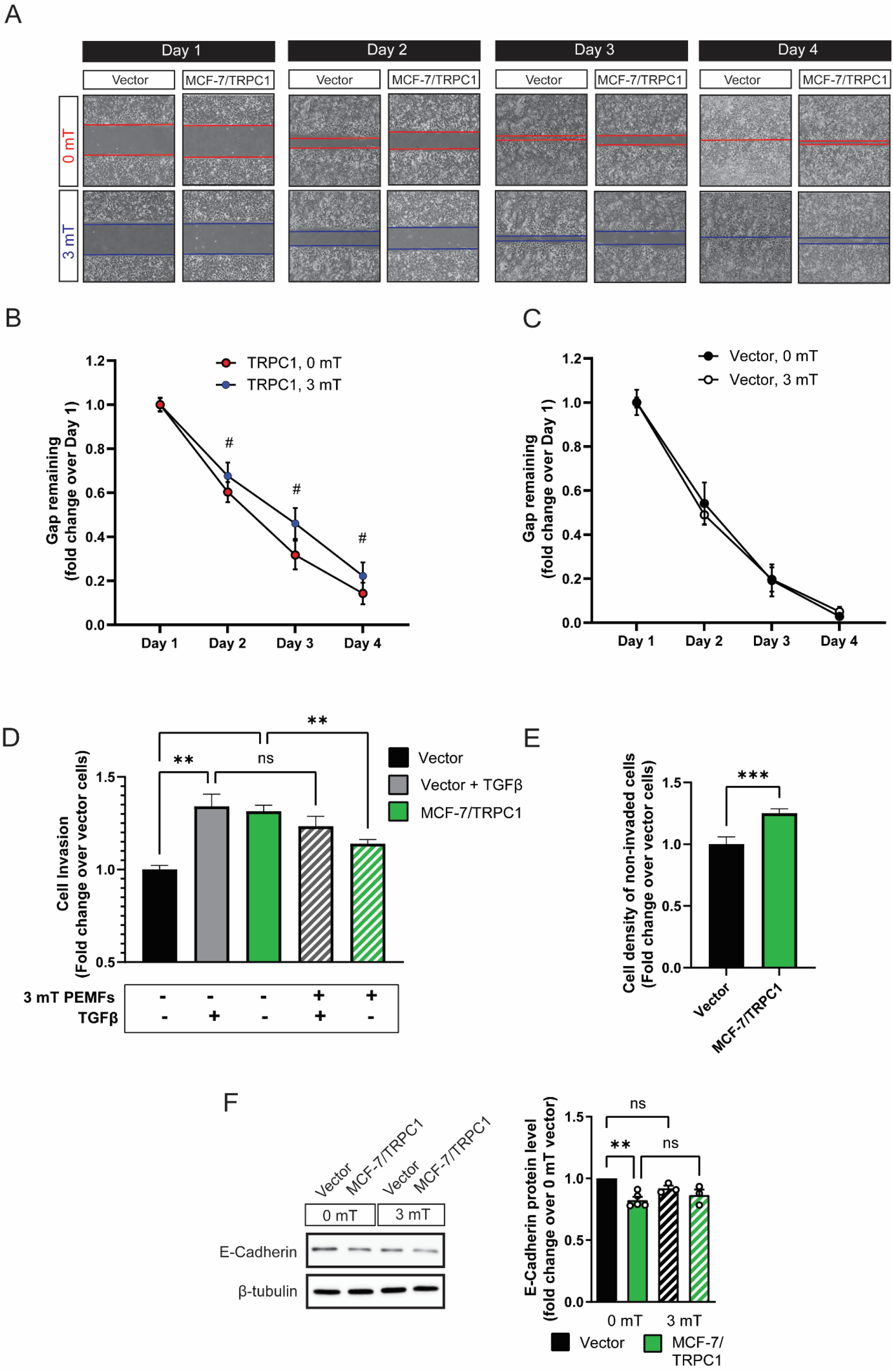
PEMF exposure attenuates migration and invasion of MCF-7/TRPC1 cells. **A)** Representative images showing the migration of vector-transfected and MCF-7/TRPC1 cells exposed to 0 or 3 mT PEMFs. Cells were plated at 30,000 cells per gap of the culture insert on day 0 and allowed to settle for 24 h before the removal of the insert. Cells were exposed to 3 mT once, twice, or thrice for 1 h daily, on days 2, 3, and 4, respectively. The corresponding bar charts represent the pooled data of **B)** MCF-7/TRPC1 (0 and 3 mT), and **C)** vector (0 and 3 mT), showing the remaining gap expressed as normalized fold change to day 1 of their respective cell line. **D)** Bar chart showing stained invasive cells at the bottom of the basal membrane expressed as fold change over vector cells. The stained cells correspond to those successfully invading the basal membrane after 48 h. Untreated vector cells serve as a control for basal cell invasion (black). The second (solid grey) and fourth (hatched grey) bar show vector cells that had been treated with TGFβ during seeding to promote invasion at plating and 24 h later. The hatched bars correspond to cells exposed to 3 mT PEMF. **E)** Analysis of cell density on the upper insert of the chamber after 48 h post-seeding. The cells were stained and lysed using the same schedule as for the invasion assay. **F)** Western analysis showing E-cadherin protein expression in vector and MCF-7/TRPC1 cells with 3 mT PEMF exposure for 1 h for 3 consecutive days. The bar chart represents fold change pooled data for E-cadherin protein expression levels normalized to 0 mT of vector cells. All results were of at least 3 independent experiments with ^*^*p* < 0.05, ^**^*p* < 0.01, ^#^*p* < 0.0001. The error bars are expressed as the standard error of the mean.

### TRPC1 overexpression increases breast cancer cell sensitivity to DOX and PEMFs

TRPC1 downregulation induces chemoresistance in ovarian cancer (19). Analogously, chronic DOX exposure reduced TRPC1 expression and promoted DOX-resistance in MCF-7 cells (Fig. 5B and 5C). Whether TRPC1 overexpression enhanced sensitivity to DOX and/or PEMF exposure was next examined. DOX was administered to MCF-7/TRPC1 cells for 4 days with or without PEMF exposure for 3 days (Fig. 8A). PEMF exposure *per se* mitigated cell growth for both MCF-7/TRPC1 (Fig. 8B; green, solid and hatched) and vector (black, solid and hatched) cells. A low dose of DOX (10 nM) produced depressions of proliferation for both MCF-7/TRPC1 (32%) and vector cells (20%), yet precluded a response to PEMF exposure in either scenario (Fig. 8B). By contrast, a higher dose of DOX (20 nM) produced strong proliferation depressions in both MCF-7/TRPC1 (69%) and vector (77%) cells that were further augmented by PEMF exposure.

**Figure 8.**
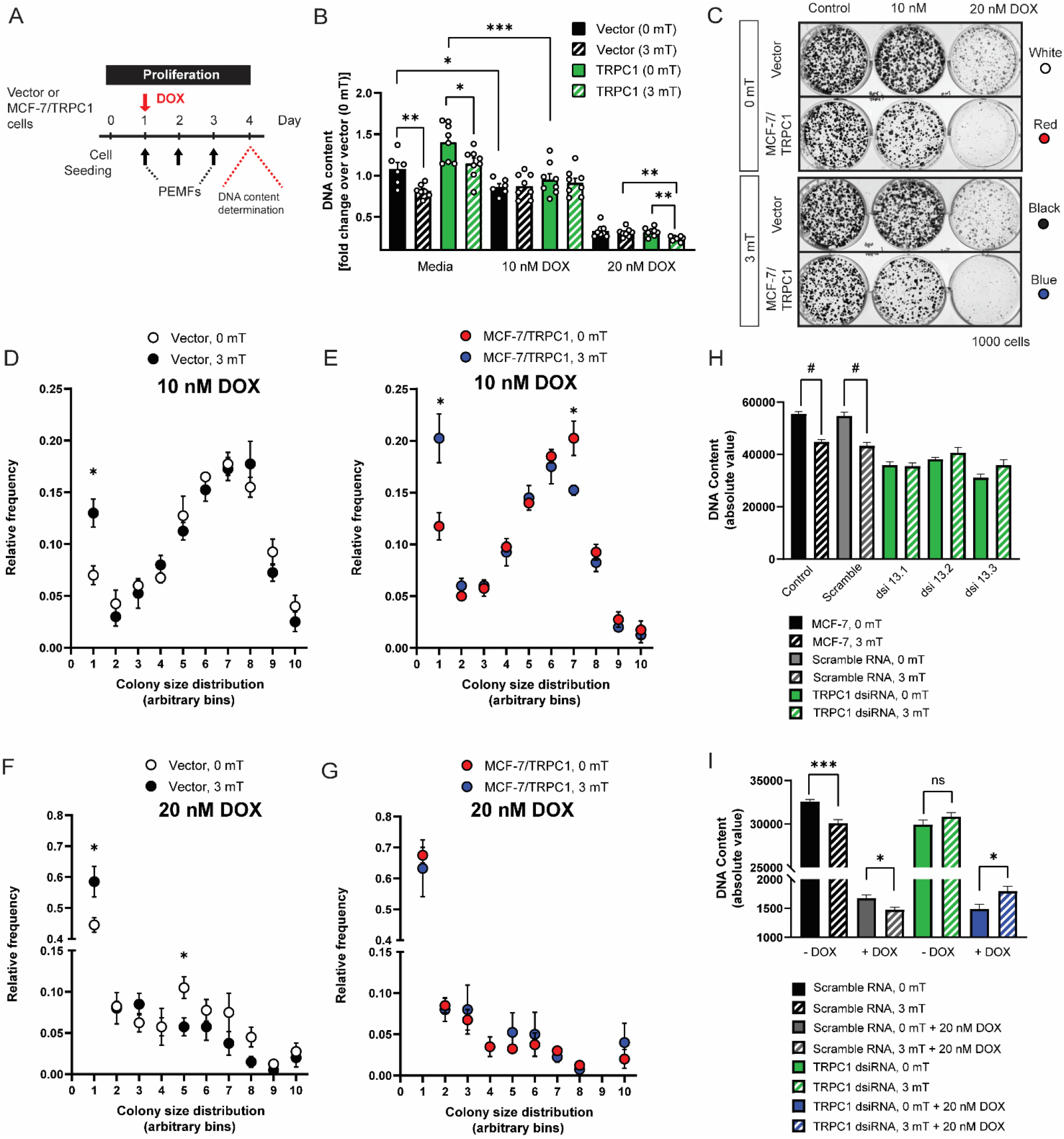
TRPC1 overexpression sensitizes breast cancer cells to doxorubicin and PEMFs. **A)** Schematic of PEMF and DOX treatment paradigms on MCF-7/TRPC1 cells for prolferation assessment using Cyquant DNA content analysis. **B)** Bar chart represents pooled data showing fold change of DNA content normalized to 0 mT vector cells. Statistical analysis was done using the two-sample *t*-test. **C)** Representative colony formation assay over 11 days of cells treated with 10 nM or 20 nM DOX, with or without 3 mT PEMF exposure. Colony size-frequency distribution histogram normalized to the total number of colonies in the presence of either **D)** 10 nM DOX on vector, **E)** 10 nM DOX on MCF-7/TRPC1, **F)** 20 nM DOX on vector, or **G)** 20 nM DOX on MCF-7/TRPC1 cells, in combination with daily PEMF exposure. The * in (D), (E), and (F) represents a statistical difference between 0 mT and 3 mT of the respective mean relative frequencies compared within the same colony size bin. **H)** Proliferation of *TRPC1*-silenced MCF-7 cells 48 h post dsiRNA transfection. Cells were transfected with three independent dsiRNA (green), including a scramble RNA (grey). Control cells (black) were left untreated but exposed to the same 0 mT (solid bars) or 3 mT (hatched bars). **I)** Combined effect of PEMF and DOX treatments on the proliferation of *TRPC1*-silenced cells. Cells were treated with 20 nM DOX and PEMF 24 h post dsiRNA transfection followed by another exposure of PEMF one day before DNA content analysis. Hatched bars represent cells exposed to 3 mT PEMF. The data for *TRPC1* dsiRNA (green and blue) was pooled data from two independent *TRPC1* dsiRNAs. The statistical analysis was generated using Multiple unpaired *t*-test for the comparison of two sample means within the same colony size. All experiments were of at least 3 independent experiments with ^*^*p* < 0.05, ^**^*p* < 0.01, ^***^*p* < 0.001, and ^#^*p* < 0.0001. The error bars represent the standard error of the means.

Colony-forming assays demonstrated an enhanced vulnerability of MCF-7/TRPC1 cells to low doses (10 nM) of DOX (Fig. 8C, middle column, second row), resulting in fewer and smaller colonies compared to vector cultures (Fig. 8C, middle column, top row). While higher doses (20 nM) of DOX further reduced colony number substantially, colony size was only modestly reduced (Fig. 8C, right column) and an effect of PEMF exposure (Fig. 8C, left column, second and bottom rows) was less obvious than that of TRPC1 overexpression *per se* (Fig. 8C, left column, 3rd and bottom rows). TRPC1 overexpression in itself was sufficient to enhance susceptibility to 10 nM DOX as depicted by the differences in the white (vector) and red (TRPC1 overexpressors) circles (Fig. 8D and 8E) in the colony size distribution plots. PEMF exposure produced a relative shift towards more numerous smaller colonies (bin 1) and fewer larger colonies (bin 7) in MCF-7/TRPC1 cells (Fig. 8E) relative to vector cells (Fig. 8D). TRPC1 overexpression appears to have made the MCF-7/TRPC1 cells hypersensitive to 20 nM DOX to the point of precluding any further PEMF-induced attenuation in colony size (Fig. 8G; blue dots), contrasting with our previous finding with naïve MCF-7 cells of PEMF-induced colony size attenuation in the presence of 20 nm DOX (Fig. 3F). On the other hand, *TRPC1*-silenced cells (Fig. 8H; green, dsiRNA 13.1, 13.2, and 13.3) exhibited reduced basal proliferation and were insensitive to PEMF exposure (hatched green), whereas untransfected (hatched black) or scramble RNA transfected cells (hatched grey) showed depressed proliferation in response to PEMF exposure. In the presence of 20 nM DOX, PEMF synergistically attenuated cell proliferation in the scramble RNA-transfected cells (Fig. 8I, solid and hatched grey), but not in *TRPC1*-silenced cells (solid and hatched blue), demonstrating that TRPC1 channel expression level establishes both DOX- and PEMF-sensitivities.

## Discussion

Initial evidence was provided of the anti-cancer attributes of an analogous pulsing magnetic field paradigm as employed in this study (8). We corroborated these earlier results by showing that identical PEMF exposure mitigated MCF-7 growth without affecting that of nonmalignant MCF10A cells and moreover, extended cancer-specific PEMF-induced cytotoxicity to include the MDA-MB-231 breast cancer cell line. These findings were further substantiated *in vivo* using the CAM model as a host for MCF-7 tumors. Consistent with our *in vitro* findings, the CAM-bearing tumors showed a significant attenuation in tumor weight and size in response to PEMF exposure (Fig. 2B).

We also provided evidence that PEMF therapy demonstrates potential to serve as a companion therapy to conventional chemotherapy. Synergism between PEMF exposure and DOX administration was explored using acute and chronic *in vitro* paradigms. Under the acute paradigm, MCF-7 cells were exposed to PEMFs over 3 successive days and once administered the *in vitro* IC_50_ dose of DOX (100 nM), whereas under the chronic paradigm MCF-7 cells were exposed to PEMFs for 10 successive days with thrice replenishment of DOX at subacute doses of 10-20 nM. Therapeutic synergism between PEMF and DOX treatments was observed in both the acute and chronic paradigms, demonstrating potentiated depressions in proliferation (Fig. 3B) and colony-growing capacity (Fig. 3D, 3E, and 3F), respectively. PEMF exposure also synergized with pemetrexed and cisplatin when tested in the acute paradigm, although with lower efficacy than with DOX (Supplementary Fig. 3A and 3B).

PEMF and DOX treatments modulate cancer viability via their mutual abilities to enhance oxidative stress (3, 5). PEMF exposure produces ROS by stimulating mitochondrial respiration (5), whereas DOX-induced ROS production is both cytoplasmic and mitochondrial in origin and arises from a process of redox cycling that instead is detrimental to mitochondrial function (3). We show that PEMF and DOX treatments synergize to augment ROS production, creating a sufficiently critical oxidative environment to induce cancer cell killing *in vitro*. The *in vivo* efficacy of PEMF and DOX treatments, separately or in combination, was demonstrated in NSG mice. The greatest reductions in patient-derived tumor size were accompanied by the largest increases in apoptosis and occurred by preceding 3 weeks of DOX chemotherapy with 2 weeks of PEMF exposure. In humans, magnetic therapy offers the advantage of being targetable to a body region inflicted with cancer for localized synergism with systemic DOX administration, potentially allowing for the lowering of chemotherapeutic dose and reducing the severity of collateral cytotoxic DOX-TRPC channel interactions, such as doxorubicin-induced cardiotoxicity (40).

### Magnetic Mitohormesis in Cancer

Mitohormesis describes a developmental process whereby low levels of oxidative stress instill mitochondrial survival adaptations by augmenting a cell’s antioxidant defenses, whereas exaggerated elevations in ROS overwhelm a cell’s existing antioxidant defenses to instead stymie cell survival (7). TRPC1 function was shown to be necessary and sufficient to confer mitochondrial responses to magnetic fields, ultimately invoking a novel process of magnetic mitohormesis (9). PEMF exposure undermines MCF-7 cell growth (Fig. 3B, 3E, and 3F) in correlation with TRPC1 expression (Fig. 5E), possibly due to the over-stimulation of this recently elucidated calcium-mitochondrial axis (Fig. 3G) (5, 9). This same PEMF protocol was better tolerated by MDA-MB-231 breast cancer cells that exhibit lower expression levels of TRPC1 (Fig. 1C, 5E). Our results are in general agreement with previous studies drawing a correlation between TRPC1 expression and the malignancy status of several forms of cancer (17, 19, 34). Magnetic-sensitivity and downstream mitochondrial activation was previously shown to be specifically correlated with TRPC1 developmental and genetic expression and function (5, 9, 27, 41), whereas the expression of other TRP channels did not show such a strong correlation (5, 27, 41) and genetic silencing of TRPM7 was unable to preclude magnetic sensitivity (5). The magnetic sensitivity conferred by TRPC1 and its relevance to mitohormetic survival mechanisms make it a valuable target for clinical exploitation in cancer treatment (5, 9, 27).

### TRPC1 Channel in Cancer

Elevated TRPC1 expression is associated with hypoxia-induced EMT in breast cancer cells (42). Here, TRPC1 overexpression was shown to increase MCF-7 proliferation and sensitivity to DOX yet, reduced migratory capacity. In a similar manner, the overexpression of miR-146b, an inflammatory modulator (43), enhanced the proliferation and chemosensitivity (cisplatin and paclitaxel) of epithelial ovarian carcinoma cells while attenuating migratory capacity (44). Therefore, under modest inflammatory conditions, such as those induced with the overexpression of TRPC1 or miR-146b, the proliferative capacities and chemosensitivities of certain cancers increase, whereas migratory capacities are diminished. Provocatively, these dichotomous proliferative and migratory reponses to inflammatory conditions may represent a point of vulnerability to be exploited in cancer treatment with PEMF-based therapies. PEMF exposure attenuated the proliferation and further slowed the migration of breast cancer cells in correlation with TRPC1 channel expression, aligning with evidence that catalytic activation of TRPC6, similarly implicated with proliferation and inflammatory responses in breast cancer, attenuated MDA MB 231 breast cancer cell viability and migratory capacity (45). Both studies further demonstrated reductions in breast cancer cell invasiveness in response to activation of either TRPC1 (PEMF exposure) (Fig. 7D) or TRPC6 (Furin inhibition) (45). These findings demonstrate the value of inducing TRPC-mediated inflammatory responses for attenuating breast cancer invasiveness and allude to a therapeutic niche for PEMF-based therapies in cancer treatment.

EMT is a multifaceted process whereby transformed cells acquire metastatic capabilities and resistance to apoptosis (36, 46). Given that small histological grade 1 breast tumors exhibit higher TRPC1 expression than larger grade 3 breast tumors (17), an elevation in TRPC1 levels may account for the propensity of small grade 1 breast tumors to undergo EMT (42, 47). In accordance, we demonstrate elevated expressions of *SLUG, SNAIL*, and *VIMENTIN* and downregulated expression of E-cadherin in TRPC1-overexpressing MCF-7 cells (Fig. 6H), consistent with metastatic induction (35). Conversely, TRPC1-silencing reduced the expressions of *SLUG* and *VIMENTIN* and upregulated E-cadherin (Fig. 6K). Elevations of TRPC1 expression are common in breast cancer (17, 42, 48) and may predispose pre-neoplastic cells towards EMT by conferring a more proliferative and invasive phenotype, but may not be required for a systemically metastatic phenotype and possibly selected against by systemic chemotherapy (*cf* Fig. 5). Indeed, negative selection by DOX against cancer cells with elevated TRPC1 expression may contribute to the commonly described chemotherapy paradox, hallmarked by the selection of cells with heightened chemoresistance (49).

Indications of cytotoxic synergies between DOX and TRPC channels exist (40). Chronic exposure of MCF-7 cells to DOX, attenuated TRPC1 expression (Fig. 5B, C) resulting in DOX-resistant cells characterized by slowed proliferation (Fig. 5D) and lost responsiveness to PEMF exposure (Supplementary Figure 4A). On the other hand, serial passaging of the DOX-resistant MCF-7/ADR cells in the absence of DOX selective pressure restored proliferative capacity (Fig. 5D) and sensitivity to PEMF exposure (Supplementary Figure 4A). While DOX treatment (100 nM) was still capable of attenuating proliferation in both MCF-7/ADR (96 nM DOX) and MCF-7/ADR (0 nM DOX), they were insensitive to PEMF exposure, suggesting irrecoverable mitochondrial damage. The clinical elaboration of PEMF-based therapies may ultimately permit the lowering of chemotherapeutic load to help avert collateral cytotoxicity (3) and paradoxical effects (49) associated with high clinical doses of DOX.

## Conclusion

TRPC1 is a mitohormetic determinant governing cellular inflammatory status and survival (5, 6, 9), whose elevated expression defines numerous cancers (13, 50). We demonstrate that the vulnerability of breast cancer to PEMF and DOX therapies is correlated with TRPC1 expression, conferring a heightened level of specificity for such TRPC1-characterized cancers (Fig. 8B to 8G). Many cancers exist near the threshold of metabolic cytotoxicity where even moderate enhancements in cellular metabolism are sufficient to cause homeostatic disequilibrium. As the TRPC1-PEMF-DOX axis exerts its actions by elevating oxidative stress, it potentially may be exploited as a therapeutic paradigm to induce cancer-specific metabolic catastrophe. The presented pulsing magnetic field paradigm in combination with systemic DOX-based chemotherapy may hence ultimately prove more selective than conventional therapies for common cancers characterized by elevated TRPC1 channel expression. Finally, given the demonstrated specificity of PEMF treatment for TRPC1 expression reported here and elsewhere (5, 9, 27, 41), in addition with its nominal effects on TRPC1 expression levels, complementation of conventional DOX-based chemotherapy by localizable PEMF therapy may help avert collateral toxicity (3) and paradoxical effects (49), by allowing the lowering of systemically-delivered chemotherapeutic dose while maintaining a unique level of specificity for TRPC1-associated cancers. The values of these possibilities merit future investigation and clinical validation.

## List of abbreviations

BCA: bicinchoninic acid
CAM: chicken chorioallantoic membrane
DCH_2_FDA: 2’,7’-dichlorodihydrofluorescein diacetate
DMEM: Dulbecco’s Modified Eagle Medium
DOX: Doxorubicin
dsiRNA: dicer-substrate short interfering RNA
EMT: epithelial-mesenchymal transition
IC_50_: half maximal inhibitory concentration
mT: milliTesla
NFAT: nuclear factor of activated T-cells
NOD-SCID: nonobese diabetic/severe combined immunodeficiency
NSG: NOD-SCID gamma
PDX: patient-derived xenograft
PEMF: pulsed electromagnetic field
PVDF: polyvinylidene difluoride
ROS: reactive oxygen species
RPMI: Roswell Park Memorial Institute
TBH: tert-butylhydroperoxide
TGFβ: Transforming Growth Factor beta
TRPC1: Transient Receptor Potential Cation Channel C Member 1

## Declarations

### Ethics approval and consent to participate

Patient samples were collected based on National Healthcare Group Domain Specific Review Board approval (2014/01088). All animal work was completed under Nanyang Technological University (NTU) Institutional Animal Care and Use Committee approval (ARF-SBS/NIE-A0141AZ, A0250AZ, A0324, and A0321).

### Consent for publication

Not applicable

### Availability of data and materials

The data supporting the conclusions of this article have been given in this article and its additional files.

### Authors’ contributions

YKT, AFO and NST conceived and designed this study. YKT, KKWC, CHHF and SR performed the cellular proliferation, ROS, RNA and protein experiments and analyses. YKT, CHHF, JLYY, and JNY generated and characterized the MCF-7 stable cell line overexpressing TRPC1. YKT, CHHF, KKWC and SR performed colony-forming assays, migration and invasion assays. RYJH and APFK established the CAM model. KKWC, SR, YKT and CHHF performed the MCF-7 cells on CAM model. CWC provided clinical human breast tumors. YSY and WRT performed the pre-clinical PDX model using human breast tumors and cell lines. YKT, KKWC, CHHF, SR, NST, AFO compiled and analyzed the data. YKT and AFO wrote the manuscript and all authors approved the final manuscript.

## Acknowledgments

The authors acknowledge Zac Goh (iHealthtech, National University of Singapore) for the design of the graphical abstract of the CAM model in Fig. 2 and the animal model in Fig. 4 We would also like to appreciate the members of the Biolonic Currents Electromagnetic Pulsing Systems Laboratory (BICEPS) for their assistance, without which this research would not be possible.

## Supplementary Figures

**Supplementary Figure 1.**
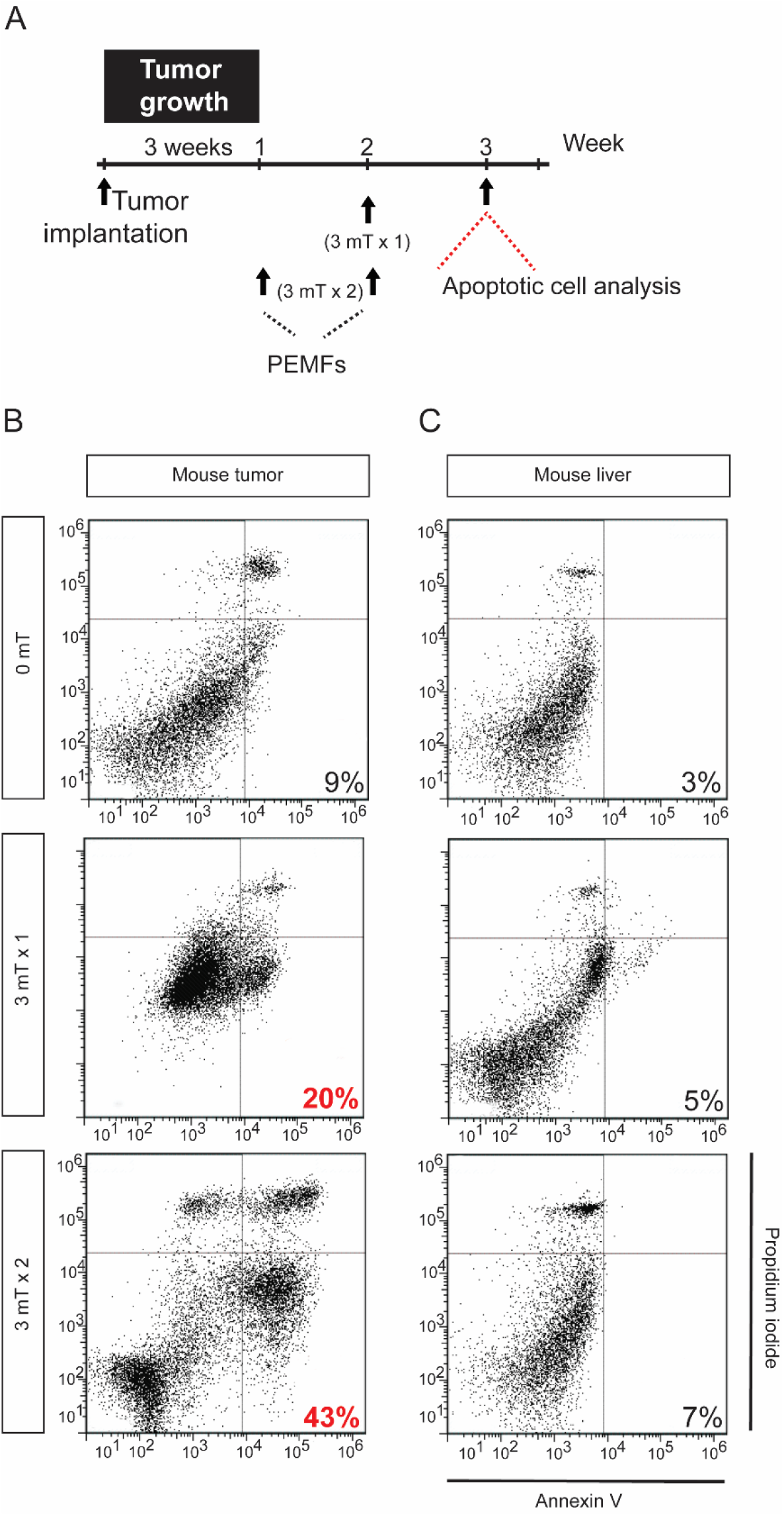
PEMFs inhibit MDA-MB-231 tumor growth without affecting liver *in vivo*. **A)** Schematic of PEMF exposure of NSG mice implanted with MDA-MB-231 tumors. Implanted tumors were allowed to grow for 3 weeks before the initiation of PEMF exposure once (3 mT x 1; 1 h, once a week) or twice (3 mT x 2; 1 h once a week for 2 weeks). Flow cytometric analysis was performed 1 week after the last PEMF exposure. Representative scatter dot-plots of **B)** MDA-MB-231 xenografts and **C)** mouse livers showing cell population of dissociated tumors sorted based on annexin and propidium iodide staining. The percentages represent the total early and late apoptotic cells.

**Supplementary Figure 2.**
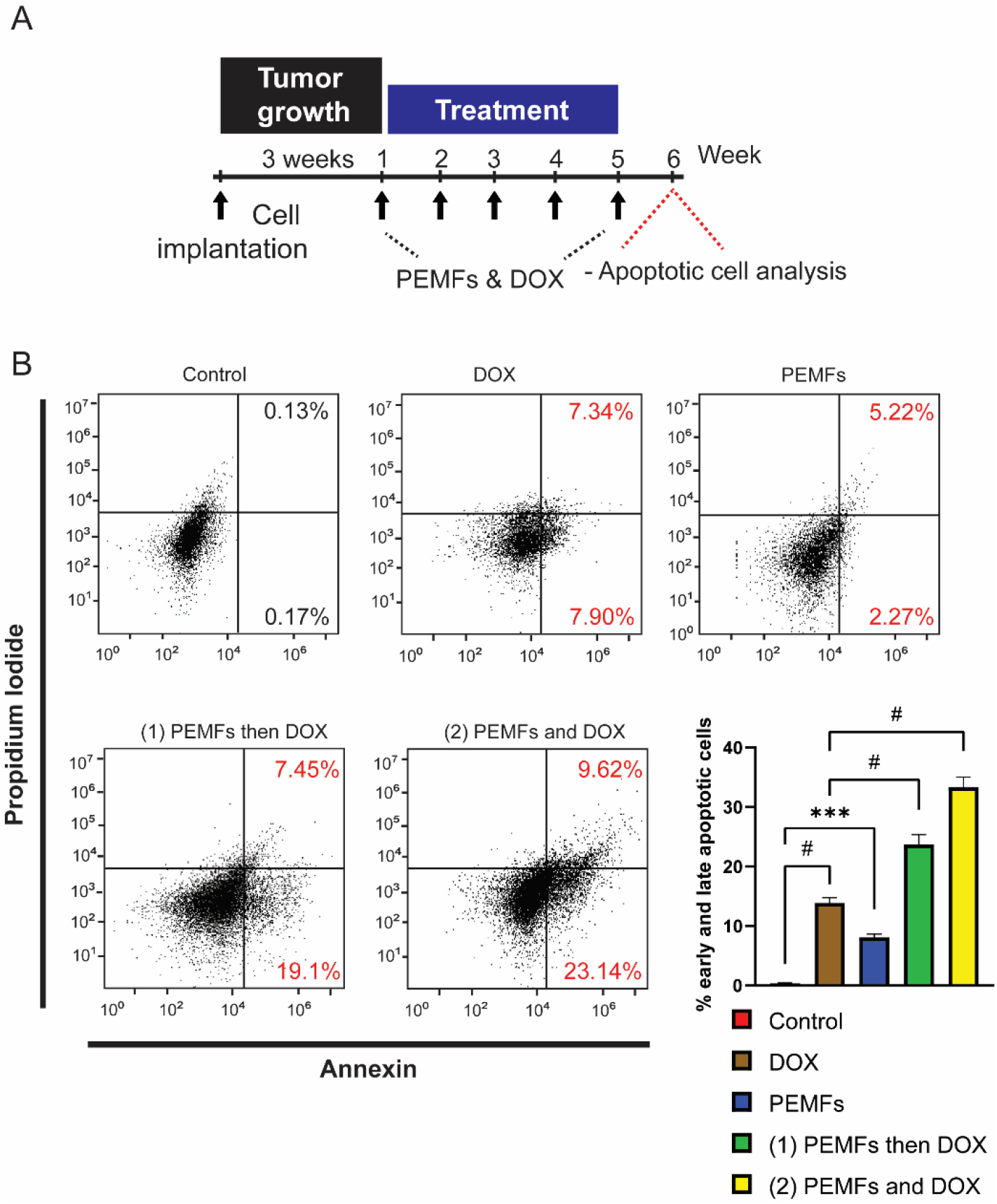
PEMFs synergize with doxorubicin to inhibit MCF-7 tumor growth *in vivo*. **A)** Schematic of weekly PEMF and DOX exposures on MCF-7 xenograft in mice. Implanted cells were allowed to grow for 3 weeks before the initiation of DOX and/or PEMF treatment. Apoptotic cell determination was performed at the end of the study. **B)** Representative scatter dot-plots showing cell population of dissociated tumors sorted based on annexin and propidium iodide staining. Bar charts represent pooled data of early and late apoptotic cell percentages analyzed using flow cytometry. N = 6 mice, with ^*\**^*p* < 0.05, ^*\*\**^*p* < 0.01, and ^*#*^*p* < 0.0001. The error bars are expressed as the standard error of the mean.

**Supplementary Figure 3.**
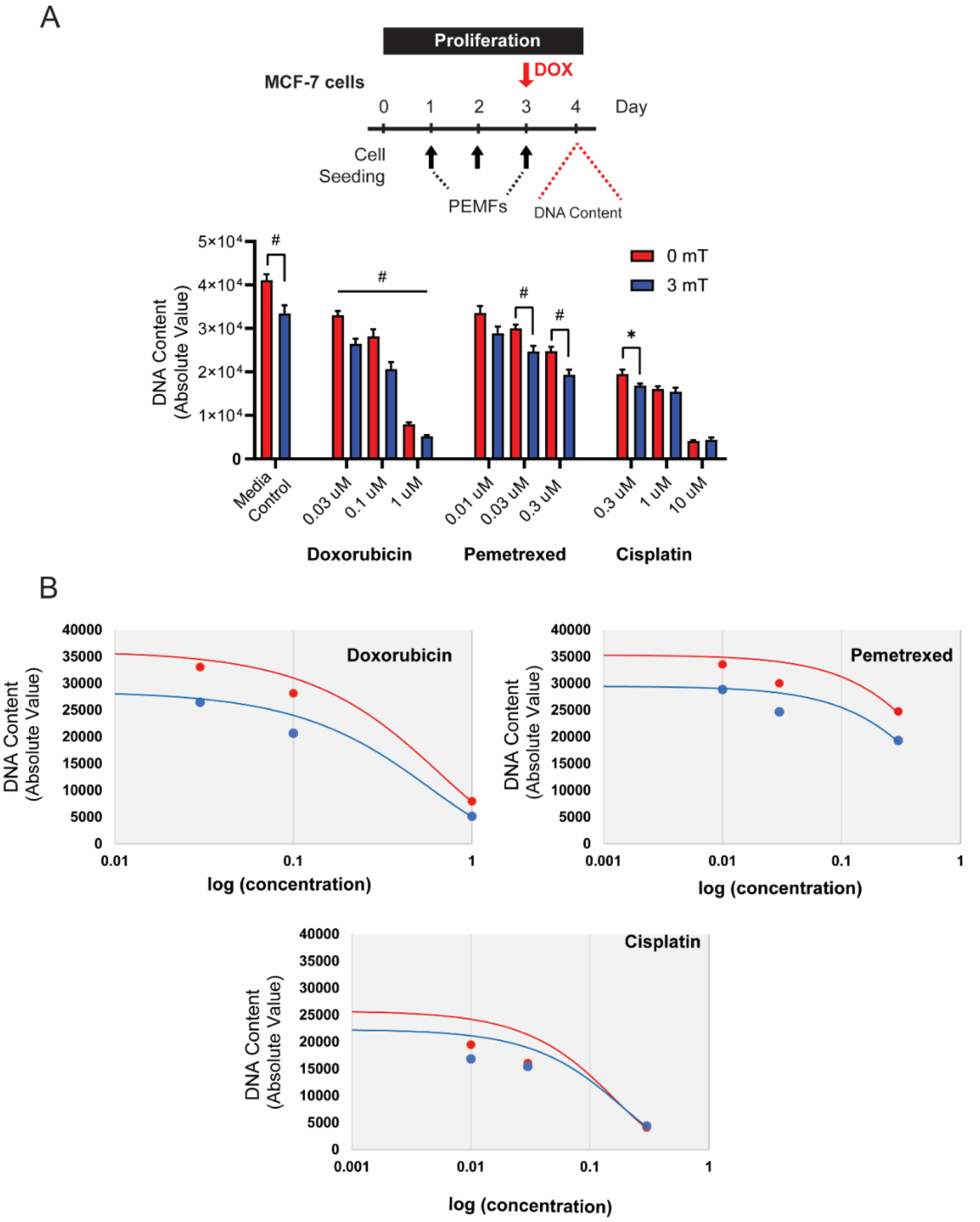
**A)** Chemo-sensitivity of MCF-7 cancer cells to Pemetrexed and Cisplatin in combination with PEMF in comparison to DOX. Cells were seeded in 96-well and exposed to 3 mT PEMF for 1 h per day for 3 days. Chemotherapeutic drugs were added (red arrow) on the final day of PEMF before the analysis of DNA content 24 h later. The corresponding bar charts show the absolute DNA content of cells in response to increasing doses of chemotherapeutic agents with or without PEMFs. **B)** Representation of a dose**-**response curve of the same data in (A), treated with DOX, Pemetrexed, and Cisplatin. The X-axis represents log (concentration) and the y-axis represents the absolute intensity of DNA content. Statistical analysis was performed using multiple unpaired *t*-tests, comparing between 0 mT and 3 mT within each concentration. All experiments were of at least 3 independent experiments with ^*^*p* < 0.05, and ^#^*p* < 0.0001. The error bars are expressed as the standard error of the mean.

**Supplementary Figure 4.**
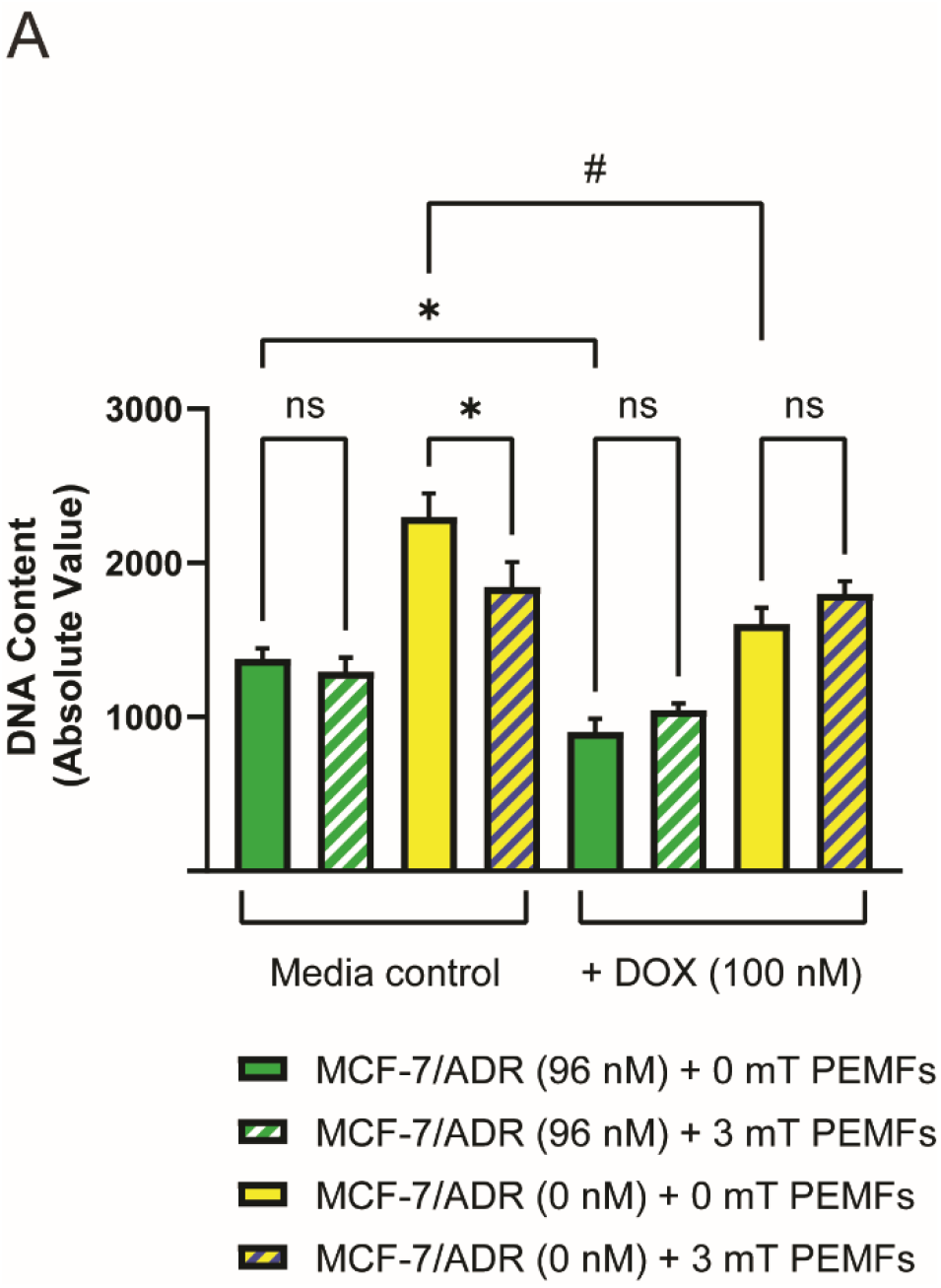
Recovery of PEMF cytotoxicity upon removal of selective pressure from DOX and reinstatement of TRPC1 expression. **A)** Cell proliferation assay using Cyquant DNA content analysis on MCF-7/ADR cell lines, in combination with DOX (100 nM) and PEMF exposure. MCF-7/ADR (96 nM) cells were maintained in 96 nM DOX while MCF-7/ADR (0 nM) were serially passaged in the absence of DOX. Cells were exposed to 3 mT PEMFs for 1 h per day for 3 days. 100 nM DOX was administered 1 h before the final PEMF exposure. Cyquant analysis was performed 24 h after the final PEMF exposure. Statistical analysis was performed using one-way ANOVA with Sidak’s multiple comparison test. All experiments were of at least 3 independent experiments with ^*^*p* < 0.05, and ^#^*p* < 0.0001. The error bars are expressed as the standard error of the mean.

## Supplementary Information

### Material and methods

#### Cell culture and pharmacology

MCF-7 (HTB-22^™^) cells were acquired from American Type Culture Collection (ATCC) and maintained in RPMI (Gibco) supplemented with 10% FBS (Hyclone) and maintained in a humidified incubator at 37°C in 5% CO_2_. MDA-MB-213 and MCF10A were acquired from Dr. Andrew Tan’s laboratory. MDA-MB-231 cells were maintained in DMEM (Gibco) and 10% FBS. MCF10A cells were maintained in growth media containing DMEM/F12 (Gibco) supplemented with 5% horse serum (Hyclone), 20 ng/ml EGF (Peprotech), 0.5 mg/ml Hydrocortisone (Sigma), 100 ng/ml cholera toxin (Sigma) and 10 ug/ml insulin (Sigma). Cells were trypsinized and passaged every 3 days using TrypLE Express reagent (Gibco). MCF-7/ADR cells resistant to 96 nM DOX were generated using a progressive incubation of cells in low 0.3 nM up to 96 nM DOX over 4 months. The concentration of DOX was doubled weekly upon cell reseeding. DOX Doxorubicin hydrochloride (DOX) (Abcam, ab120629) was reconstituted in DMSO to make a stock concentration of 25 mM and stored at −80°C. Subsequent dilutions of DOX were made in distilled water to keep DMSO concentration below 0.01%. For the characterization of TRPC1 protein levels, PC3 and LNCaP cells were maintained in RPMI media containing 10% FBS. No cell culture antibiotics were used throughout the experiments.

#### Cell count and DNA content analysis

For cell enumeration using trypan blue exclusion assay, MCF-7, MDA-MB-231, or MCF10A cells were seeded at 6000 cells/cm^2^ per well of a 6-well plate. For MCF10A cells, they were plated in growth media without EGF. Cell counting was performed using 3 wells of a 6-well plate for technical replication. For DNA content analysis using Cyquant cell proliferation assay (Invitrogen), cells were seeded at 4000 cells per well and performed with 8 technical replicates in a 96-well plate. Seeded cells were left for 24 h before treatment with DOX or exposed to PEMFs. Cyquant stained DNA was measured using at 480/520 nm using Cytation 5 microplate reader (BIOTEK).

#### Clonogenic assay and quantification of colonies

*In vitro* clonogenic assay was performed using crystal violet staining (22). Briefly, MCF-7 cells were seeded either at 100 or 1000 cells per well of a 6-well plate. The cells were treated with DOX on Day 1, 4, and 7 in RPMI supplemented with 10% FBS. 3 mT PEMFs stimulation was administered for 1 h from Day 1 to Day 10. On Day 11, the cells were rinsed in PBS and stained with crystal violet stain consisting of 0.5% crystal violet and 25% glutaraldehyde (Sigma Aldrich) in distilled water for 3 h. Stained colonies were rinsed with 2 changes of tap water and left to dry. Images of the colonies were taken using Chemidoc Imaging System (BIORAD) under the Coomassie Blue Stain filter setting. The number of colonies and colony size per well was estimated using the ImageJ Analyze particle option using 3 to 3500-pixel unit with a circularity of 0.2 to 1. The mean survival factor was determined as the number of surviving cells over the number of cells plated and normalized to the survival factor of the control group expressed as fold change. The colony size relative frequency was determined by binning colonies into 10 bins, according to their relative size from the smallest (1) to the largest (10) colonies after normalizing to the total number of cells.

#### Reactive oxygen species analysis using DCH_2_FDA

Cells were seeded in 96-well clear bottom black well (Costar) at a density of 10,000 cells per well at 8 replicates per condition and left to settle for 24 h before commencing the experiment. Cells were rinsed with warm phenol-free and serum-free (PFSF) RPMI (GIBCO) and incubated with PFSF RPMI containing 10 uM DCH_2_FDA (Invitrogen) for 30 min in a standard tissue culture incubator. The dye was then rinsed out using warm PFSF RPMI and treated with tert-Butyl hydroperoxide (TBH, 1 mM; Sigma Aldrich) or 50 uM DOX (Abcam) in PFSF RPMI. Cells were immediately exposed to 10 min of 0 mT or 3 mT PEMFs exposure before proceeding to ROS determination using a Cytation 5 microplate reader (BIOTEK) at Ex/EM: 492/520 nm for 20 min with the temperature set at 37°C.

#### Western Analysis

Cell lysates were prepared in ice-cold radioimmunoprecipitation assay (RIPA) buffer containing 150 mM NaCl, 1% Triton X-100, 0.5% sodium deoxycholate, 0.1% SDS and 50 mM Tris (pH 8.0) supplemented with protease and phosphatase inhibitors (PhosphoSTOP, Roche). Cells were lysed for 20 min and centrifuged for 10 min at 13,500 rpm. The protein concentration of the soluble fractions was determined using a BCA reagent (Thermo Fisher Scientific). 25 - 50 ug of total protein was resolved using 10% or 12% denaturing polyacrylamide gel electrophoresis and transferred to PVDF membrane (Immobilon-P, PVDF). Proteins on PVDF membranes were blocked using 5% low-fat milk in TBST containing 0.1% Tween-20 and incubated with the primary antibody in SuperBlock (TBS, Thermo Fisher Scientific) overnight at 4 °C. The primary antibodies used were: TRPC1 (1:300; Santa Cruz), Cyclin D1 (CD1, 1:300; Santa Cruz), Caspase 3 (1:300; Santa Cruz), GFP (1:1000; Proteintech), β-actin (1:10,000; Proteintech), α-tubulin (1:5000; Proteintech). The membranes were washed in TBST. Anti-rabbit or anti-mouse antibody conjugated to horseradish peroxidase (HRP) were diluted (1:3000, Thermo Fisher Scientific) in 5% milk in TBST and were incubated with the membranes for 1 h at room temperature. The membranes were incubated in WestPico or WestFemto chemiluminescent substrate (Thermo Fisher Scientific), detected and analyzed using LI-COR Image Studio.

#### Laser confocal imaging

For the visualization of GFP and Vimentin abundance in TRPC1 overexpressing MCF-7 cells, the cells were seeded onto coverslips at a density of 100,000 cells per well of a 6-well plate (NUNC). 24 h post-seeding, the cells were rinsed with PBS and fixed in 4% paraformaldehyde for 20 min. For the direct visualization of the expression of GFP in vector-only and MCF-7/TRPC1 cells, the cells on coverslips were mounted onto glass slides using ProLong Gold Antifade Mountant (Thermo Fisher Scientific). The cells were then analyzed using the Olympus FV1000 confocal laser scanning microscope. For the visualization of *VIMENTIN*, the cells were permeabilized with 0.1% Triton in PBS for 10 mins after fixation. The cells were then blocked in SuperBlock TBS (Thermo Fisher Scientific) followed by Vimentin antibody (Santa Cruz, 1:100) incubation overnight, followed by secondary Alexa Fluor 594 antibody (1:500, Thermo Fisher Scientific) for 1 h at room temperature. Washes between steps were done with PBS with 0.1% Tween (Sigma Aldrich). Nuclei of cells were co-stained with DAPI (Sigma Aldrich) for 10 min. Cells were finally mounted and visualized using a laser confocal microscope. For the quantitative analysis of Vimentin abundance, the total absolute intensity per view was normalized to the number of nuclei to yield a mean protein intensity per cell. The average of the mean protein intensity per cell (at least 10 cells per view) from multiple replicates were used to compute and compare the abundance of Vimentin protein between vector-only and MCF-7/TRPC1 cells.

#### Real-time qPCR and TRPC1 silencing

Quantitative reverse-transcription polymerase chain reaction (RT-qPCR) was carried out using the SYBR green-based detection workflow. Briefly, total RNA was harvested from MCF-7 cells using Trizol reagent (Thermo Fisher Scientific) and 0.5 ug of RNA was reverse transcribed to cDNA using iScript cDNA Synthesis kit (Bio-Rad). Quantification of gene transcript expression was performed using SSoAdvanced Universal SYBR Green (Bio-Rad) on the CFZ Touch Real-Time PCR Detection System (Bio-Rad). Relative transcript expression was determined using the 2^-ΔΔCt^ method, normalized to β-actin transcript levels. The qPCR primers used were: *TRPC1*, F: 5’-AAG CTT TTC TTG CTG GCG TG, R: 5’-ATC TGC AGA CTG ACA ACC GT; *SNAIL*, F: 5’-CGA GTG GTT CTT CTG CGC TA, R: 5’-CTG CTG GAA GGT AAA CTC TGG A; *SLUG*, F: 5’-TAG AAC TCA CAC GGG GGA GAA G, R: 5’-ATT GCG TCA CTC AGT GTG CT; *VIMENTIN* F: 5’-AAG GCG AGG AGA GCA GGA TT, R: 5’-AGG TCA TCG TGA TGC TGA GA; and β-actin, 5’-AGA AGA TGA CCC AGA TCA TGT TTG A, R: 5’-AGC ACA GCC TGG ATA GCA AC.

For TRPC1 silencing in MCF-7 cells, two pre-designed dicer-substrate short interfering RNAs (dsiRNA, IDT) were used to knock down the expression of TRPC1. Both dsiRNAs targeted the coding-sequence of TRPC1 (NM_001251845). Transfection of dsiRNA was performed using Lipofectamine 3000 reagent (Invitrogen) as per manufacturer’s protocol. TRPC1-silenced cells were validated using qPCR 48 h post dsiRNA transfection using primers against *TPRC1, SNAIL, SLUG* and *VIMENTIN* as indicated above, relative to cells transfected with scramble dsiRNA.

#### Migration Assay

MCF-7 cells at a density of 30,000 cells in 120 ul RPMI supplemented with 10% FBS were seeded into each gap of a 4-well 3.5 mm culture dish insert (ibidi). The cells were left to adhere for 24 h before the removal of the insert and the addition of RPMI media containing 10% FBS to a total volume of 2 ml per dish. Closure of the gaps was captured using light microscopy on all four limbs of the insert, taken every 24 h. The average of 16 gap distances was considered from the 4 limbs with 4 readings arising from each limb. The images of the gap distances were analyzed using ImageJ.

#### Invasion Assay

Invasion assay was performed using the CytoSelect 24-well Cell Invasion Assay kit (Cell Biolabs, Inc.) according to the manufacturer’s protocol. Briefly, 300,000 cells were seeded in the cell culture insert after the rehydration of the basal membrane in FBS-free RPMI media. The lower well of the invasion plate was filled with RPMI media supplemented with 10% to promote the invasion of cells through the basal membrane. 20 ng/ml TGFβ was added to selected conditions in the cell culture insert to stimulate cell invasion. The setup was incubated for 48 h in a standard tissue culture incubator before the extraction and staining of the invaded cells from the basal membrane. The lysates from the extracted cells were analyzed at OD 560 using Cytation 5 microplate reader (BIOTEK).

#### Generation of plasmid and stable cell line

GFP-TRPC1 plasmid was generated by PCR amplification of full-length human TRPC1 cDNA (Accession: NM_001251845.2; 2382 base pairs) and directionally subcloned into the pEGFP-C1 vector. Transfection of plasmids in MCF-7 cells was carried out using Lipofectamine 3000 reagent (Invitrogen). 48 h after plasmid transfection, stable cells were selected in RPMI containing 750 ug/ml Geneticin (Invitrogen), 10% FBS, and 1% Pen/Strep (Gibco) in 5% CO_2_ at 37 °C. GFP vector and GFP-TRPC1 cells were enriched for GFP positive cells using Beckman Coulter Moflo Astrios cell sorter. Stables cells were subsequently maintained in complete RPMI media containing 500 ug/ml Geneticin. The overexpression of GFP-TRPC1 in the stable cells was characterized using qPCR, immunofluorescence, and western analysis. GFP stable cells are referred to as vector-only cells while GFP-TRPC1 overexpression stables cells are referred to as MCF-7/TRPC1 cells in the manuscript.

#### Apoptotic assay

For apoptotic cell determination, the tumors were dissociated to single cells using the MACS Tumor Dissociation Kit in combination with the gentleMACS Dissociator (Miltenyi Biotec) as according to the manufacturer’s protocol. After dissociation, the cells were filtered through a 30 um MACS SmartStrainer. Cells were pelleted from the filtrate at 300 g x 7 min and resuspended in 400 ul Binding Buffer. The cells were incubated with Annexin V FITC and Propidium Iodide (Sigma Aldrich) for 15 min in the dark at room temperature. After incubation, the cells were pelleted and resuspended in 100 ul Binding Buffer for analysis by flow cytometry using BD Accuri C6 cytometer (BD Biosciences, CA, USA).

#### Statistical analysis

All statistics were carried out using GraphPad Prism (Version 8) software. One-way analysis of variance (ANOVA) was used to compare the values between two or more groups supported by multiple comparisons. This was followed by Bonferroni’s posthoc test. For the comparison between two independent samples, the Student’s *t*-test was performed.

## References

1. Coughlin SS, Ekwueme DU. Breast cancer as a global health concern. Cancer Epidemiol. 2009;33(5):315–8.

2. DeSantis CE, Ma J, Gaudet MM, Newman LA, Miller KD, Goding Sauer A, et al. Breast cancer statistics, 2019. CA Cancer J Clin. 2019;69(6):438–51.

3. McGowan JV, Chung R, Maulik A, Piotrowska I, Walker JM, Yellon DM. Anthracycline Chemotherapy and Cardiotoxicity. Cardiovasc Drugs Ther. 2017;31(1):63–75.

4. Kalyanaraman B, Cheng G, Hardy M, Ouari O, Bennett B, Zielonka J. Teaching the basics of reactive oxygen species and their relevance to cancer biology: Mitochondrial reactive oxygen species detection, redox signaling, and targeted therapies. Redox Biol. 2018;15:347–62.

5. Yap JLY, Tai YK, Frohlich J, Fong CHH, Yin JN, Foo ZL, et al. Ambient and supplemental magnetic fields promote myogenesis via a TRPC1-mitochondrial axis: evidence of a magnetic mitohormetic mechanism. FASEB J. 2019;33(11):12853–72.

6. Tai YK, Ng C, Purnamawati K, Yap JLY, Yin JN, Wong C, et al. Magnetic fields modulate metabolism and gut microbiome in correlation with Pgc-1alpha expression: Follow-up to an in vitro magnetic mitohormetic study. FASEB J. 2020;34(8):11143–67.

7. Ristow M, Schmeisser K. Mitohormesis: Promoting Health and Lifespan by Increased Levels of Reactive Oxygen Species (ROS). Dose Response. 2014;12(2):288–341.

8. Crocetti S, Beyer C, Schade G, Egli M, Frohlich J, Franco-Obregon A. Low intensity and frequency pulsed electromagnetic fields selectively impair breast cancer cell viability. PLoS One. 2013;8(9):e72944.

9. Kurth F, Tai YK, Parate D, van Oostrum M, Schmid YRF, Toh SJ, et al. Cell-Derived Vesicles as TRPC1 Channel Delivery Systems for the Recovery of Cellular Respiratory and Proliferative Capacities. Adv Biosyst. 2020;4(11):e2000146.

10. Boyman L, Karbowski M, Lederer WJ. Regulation of Mitochondrial ATP Production: Ca(2+) Signaling and Quality Control. Trends Mol Med. 2020;26(1):21–39.

11. Gervasio OL, Whitehead NP, Yeung EW, Phillips WD, Allen DG. TRPC1 binds to caveolin-3 and is regulated by Src kinase - role in Duchenne muscular dystrophy. J Cell Sci. 2008;121(Pt 13):2246–55.

12. Yang D, Kim J. Emerging role of transient receptor potential (TRP) channels in cancer progression. BMB Rep. 2020;53(3):125–32.

13. Chinigo G, Fiorio Pla A, Gkika D. TRP Channels and Small GTPases Interplay in the Main Hallmarks of Metastatic Cancer. Front Pharmacol. 2020;11:581455.

14. Hwang JA, Hwang MK, Jang Y, Lee EJ, Kim JE, Oh MH, et al. 20-O-beta-d-glucopyranosyl-20(S)-protopanaxadiol, a metabolite of ginseng, inhibits colon cancer growth by targeting TRPC channel-mediated calcium influx. J Nutr Biochem. 2013;24(6):1096–104.

15. Selli C, Erac Y, Tosun M. Simultaneous measurement of cytosolic and mitochondrial calcium levels: observations in TRPC1-silenced hepatocellular carcinoma cells. J Pharmacol Toxicol Methods. 2015;72:29–34.

16. Jang Y, Lee Y, Kim SM, Yang YD, Jung J, Oh U. Quantitative analysis of TRP channel genes in mouse organs. Arch Pharm Res. 2012;35(10):1823–30.

17. Dhennin-Duthille I, Gautier M, Faouzi M, Guilbert A, Brevet M, Vaudry D, et al. High expression of transient receptor potential channels in human breast cancer epithelial cells and tissues: correlation with pathological parameters. Cell Physiol Biochem. 2011;28(5):813–22.

18. Kim YA, Cho DY, Przytycka TM. Understanding Genotype-Phenotype Effects in Cancer via Network Approaches. PLoS Comput Biol. 2016;12(3):e1004747.

19. Liu X, Zou J, Su J, Lu Y, Zhang J, Li L, et al. Downregulation of transient receptor potential cation channel, subfamily C, member 1 contributes to drug resistance and high histological grade in ovarian cancer. Int J Oncol. 2016;48(1):243–52.

20. Yang B, Wolfenson H, Chung VY, Nakazawa N, Liu S, Hu J, et al. Stopping transformed cancer cell growth by rigidity sensing. Nat Mater. 2020;19(2):239–50.

21. Ito R, Takahashi T, Katano I, Ito M. Current advances in humanized mouse models. Cell Mol Immunol. 2012;9(3):208–14.

22. Franken NA, Rodermond HM, Stap J, Haveman J, van Bree C. Clonogenic assay of cells in vitro. Nat Protoc. 2006;1(5):2315–9.

23. Tsou SH, Chen TM, Hsiao HT, Chen YH. A critical dose of doxorubicin is required to alter the gene expression profiles in MCF-7 cells acquiring multidrug resistance. PLoS One. 2015;10(1):e0116747.

24. Buckner CA, Buckner AL, Koren SA, Persinger MA, Lafrenie RM. Exposure to a specific time-varying electromagnetic field inhibits cell proliferation via cAMP and ERK signaling in cancer cells. Bioelectromagnetics. 2018;39(3):217–30.

25. Osera C, Amadio M, Falone S, Fassina L, Magenes G, Amicarelli F, et al. Pre-exposure of neuroblastoma cell line to pulsed electromagnetic field prevents H2 O2 - induced ROS production by increasing MnSOD activity. Bioelectromagnetics. 2015;36(3):219–32.

26. Celik C, Franco-Obregon A, Lee EH, Hui JH, Yang Z. Directionalities of magnetic fields and topographic scaffolds synergise to enhance MSC chondrogenesis. Acta Biomater. 2021;119:169–83.

27. Madanagopal TT, Tai YK, Lim SH, Fong CH, Cao T, Rosa V, et al. Pulsed electromagnetic fields synergize with graphene to enhance dental pulp stem cell-derived neurogenesis by selectively targeting TRPC1 channels. Eur Cell Mater. 2021;41:216–32.

28. Tarpey MD, Amorese AJ, Balestrieri NP, Fisher-Wellman KH, Spangenburg EE. Doxorubicin causes lesions in the electron transport system of skeletal muscle mitochondria that are associated with a loss of contractile function. J Biol Chem. 20 19;294(51):19709–22.

29. Monteith GR, Prevarskaya N, Roberts-Thomson SJ. The calcium-cancer signalling nexus. Nat Rev Cancer. 2017;17(6):367–80.

30. Villalobos C, Hernandez-Morales M, Gutierrez LG, Nunez L. TRPC1 and ORAI1 channels in colon cancer. Cell Calcium. 2019;81:59–66.

31. Denard B, Lee C, Ye J. Doxorubicin blocks proliferation of cancer cells through proteolytic activation of CREB3L1. Elife. 2012;1:e00090.

32. Nagaraja GM, Othman M, Fox BP, Alsaber R, Pellegrino CM, Zeng Y, et al. Gene expression signatures and biomarkers of noninvasive and invasive breast cancer cells: comprehensive profiles by representational difference analysis, microarrays and proteomics. Oncogene. 2006;25(16):2328–38.

33. Liu YL, Chou CK, Kim M, Vasisht R, Kuo YA, Ang P, et al. Assessing metastatic potential of breast cancer cells based on EGFR dynamics. Sci Rep. 2019;9(1):3395.

34. Pigozzi D, Ducret T, Tajeddine N, Gala JL, Tombal B, Gailly P. Calcium store contents control the expression of TRPC1, TRPC3 and TRPV6 proteins in LNCaP prostate cancer cell line. Cell Calcium. 2006;39(5):401–15.

35. Vuoriluoto K, Haugen H, Kiviluoto S, Mpindi JP, Nevo J, Gjerdrum C, et al. Vimentin regulates EMT induction by Slug and oncogenic H-Ras and migration by governing Axl expression in breast cancer. Oncogene. 2011;30(12):1436–48.

36. Thiery JP, Acloque H, Huang RY, Nieto MA. Epithelial-mesenchymal transitions in development and disease. Cell. 2009;139(5):871–90.

37. Tavakolian S, Goudarzi H, Faghihloo E. E-cadherin, Snail, ZEB-1, DNMT1, DNMT3A and DNMT3B expression in normal and breast cancer tissues. Acta Biochim Pol. 2019;66(4):409–14.

38. Na TY, Schecterson L, Mendonsa AM, Gumbiner BM. The functional activity of E-cadherin controls tumor cell metastasis at multiple steps. Proc Natl Acad Sci U S A. 2020;117(11):5931–7.

39. Canel M, Serrels A, Frame MC, Brunton VG. E-cadherin-integrin crosstalk in cancer invasion and metastasis. J Cell Sci. 2013;126(Pt 2):393–401.

40. Chen RC, Sun GB, Ye JX, Wang J, Zhang MD, Sun XB. Salvianolic acid B attenuates doxorubicin-induced ER stress by inhibiting TRPC3 and TRPC6 mediated Ca(2+) overload in rat cardiomyocytes. Toxicol Lett. 2017;276:21–30.

41. Parate D, Franco-Obregon A, Frohlich J, Beyer C, Abbas AA, Kamarul T, et al. Enhancement of mesenchymal stem cell chondrogenesis with short-term low intensity pulsed electromagnetic fields. Sci Rep. 2017;7(1):9421.

42. Azimi I, Milevskiy MJG, Kaemmerer E, Turner D, Yapa K, Brown MA, et al. TRPC1 is a differential regulator of hypoxia-mediated events and Akt signalling in PTEN-deficient breast cancer cells. J Cell Sci. 2017;130(14):2292–305.

43. Zhang L, Dong L, Tang Y, Li M, Zhang M. MiR-146b protects against the inflammation injury in pediatric pneumonia through MyD88/NF-kappaB signaling pathway. Infect Dis (Lond). 2020;52(1):23–32.

44. Yan M, Yang X, Shen R, Wu C, Wang H, Ye Q, et al. miR-146b promotes cell proliferation and increases chemosensitivity, but attenuates cell migration and invasion via FBXL10 in ovarian cancer. Cell Death Dis. 2018;9(11):1123.

45. López JJS, G.; Cantonero, C.; Soulet, F.; Descarpentrie, J.; Smani, T.; Badiola, I.; Pernot, S.; Evrard, S.; Rosado, J.A.; Khatib, A.-M. Furin Prodomain ppFurin Enhances Ca2+ Entry Through Orai and TRPC6 Channels’ Activation in Breast Cancer Cells. Cancers (Basel). 2021;13(7):1670.

46. Iwatsuki M, Mimori K, Yokobori T, Ishi H, Beppu T, Nakamori S, et al. Epithelial-mesenchymal transition in cancer development and its clinical significance. Cancer Sci. 2010;101(2):293–9.

47. Savci-Heijink CD, Halfwerk H, Hooijer GKJ, Koster J, Horlings HM, Meijer SL, et al. Epithelial-to-mesenchymal transition status of primary breast carcinomas and its correlation with metastatic behavior. Breast Cancer Res Treat. 2019;174(3):649–59.

48. Zhang LY, Zhang YQ, Zeng YZ, Zhu JL, Chen H, Wei XL, et al. TRPC1 inhibits the proliferation and migration of estrogen receptor-positive Breast cancer and gives a better prognosis by inhibiting the PI3K/AKT pathway. Breast Cancer Res Treat. 2020;182(1):21–33.

49. D’Alterio C, Scala S, Sozzi G, Roz L, Bertolini G. Paradoxical effects of chemotherapy on tumor relapse and metastasis promotion. Semin Cancer Biol. 2020;60:351–61.

50. Elzamzamy OM, Penner R, Hazlehurst LA. The Role of TRPC1 in Modulating Cancer Progression. Cells. 2020;9(2).

